# Epigenetic phase variation in the gut microbiome enhances bacterial adaptation

**DOI:** 10.1101/2025.01.11.632565

**Authors:** Mi Ni, Yu Fan, Yujie Liu, Yangmei Li, Wanjin Qiao, Lauren E. Davey, Xue-Song Zhang, Magdalena Ksiezarek, Edward A. Mead, Alan Tourancheau, Wenyan Jiang, Martin J. Blaser, Raphael H. Valdivia, Gang Fang

## Abstract

The human gut microbiome within the gastrointestinal tract continuously adapts to variations in diet, medications, and host physiology. A strategy for bacterial genetic adaptation is epigenetic phase variation (ePV) mediated by bacterial DNA methylation, which can regulate gene expression, enhance clonal heterogeneity, and enable a single bacterial strain to exhibit variable phenotypic states. Genome-wide and site-specific ePVs have been characterized in human pathogens’ antigenic variation and virulence factor production. However, the role of ePV in facilitating adaptation within the human microbiome remains poorly understood. Here, we comprehensively cataloged genome-wide and site-specific ePV in human infant and adult gut microbiomes. First, using long-read metagenomic sequencing, we detected genome-wide ePV mediated by complex structural variations of DNA methyltransferases, highlighting those associated with antibiotics or fecal microbiota transplantation. Second, we analyzed a collection of public short-read metagenomic sequencing datasets, uncovering a great prevalence of genome-wide ePV in the human gut microbiome. Third, we quantitatively detected site-specific ePVs using single-molecule methylation analysis to identify dynamic variation associated with antibiotic treatment or probiotic engraftment. Finally, we performed an in-depth assessment of an *Akkermansia muciniphila* isolate from an infant, highlighting that ePVs can regulate gene expression and enhance the bacterial adaptive capacity by employing a bet-hedging strategy to increase tolerance to differing antibiotics. Our findings indicate that epigenetic modifications are a common strategy used by gut bacteria to adapt to the fluctuating environment.

## Introduction

The human gut microbiome is continuously subjected to a myriad of changes including those due to changes in diet, medications, and physiological states.^1–3^ In response to these alterations, the gut microbiome shows a remarkable adaptive capacity.^4–6^ Characterizing this adaptive capacity is crucial for understanding the dynamic relationship between the gut microbiome and host physiology, especially in the context of human health and disease. While many microbiome studies are focused on profiles of microbial species and the relevant functions across different human diseases,^4,5,7–11^ great progress has also been made with regard to the molecular mechanisms that underpin the microbiome’s adaptive capacity including point mutations,^12–14^ copy number variation,^15,16^ structural variation (SV),^17,18^ gene amplification,^19,20^ invertible promoters^21^ and intragenic inversions.^22^ These genomic variations provide a single bacterial strain with the plasticity of expressing alternative, heritable, and reversible phenotypic states, including phase variation.^23,24^ This mechanism increases the chances of bacterial survival and adaptation by producing a genetically diverse population that can withstand multiple stresses.^21,22^

Another mechanism of bacterial adaptation involves DNA methylation, which can regulate gene expression, enhance clonal heterogeneity and mediate epigenetic phase variation (ePV, intra-strain epigenetic variation that leads to phenotypic differences, **Fig. 1A**).^25–28^ Bacterial DNA contains three primary types of DNA methylation: N6-methyladenine (6mA), N4-methylcytosine (4mC), and 5-methylcytosine (5mC).^29^ These DNA modifications are catalyzed by DNA methyltransferases (MTases) that are either part of restriction-modification systems or orphan enzymes without cognate restriction genes.^25,29^ ePVs driven by DNA methylation have been characterized in multiple human pathogens, where it facilitates processes such as antigenic variation and virulence factor production,^30–32^ which can impact how the bacterium interacts with its environment and host.^28,33–35^

**Figure 1.**
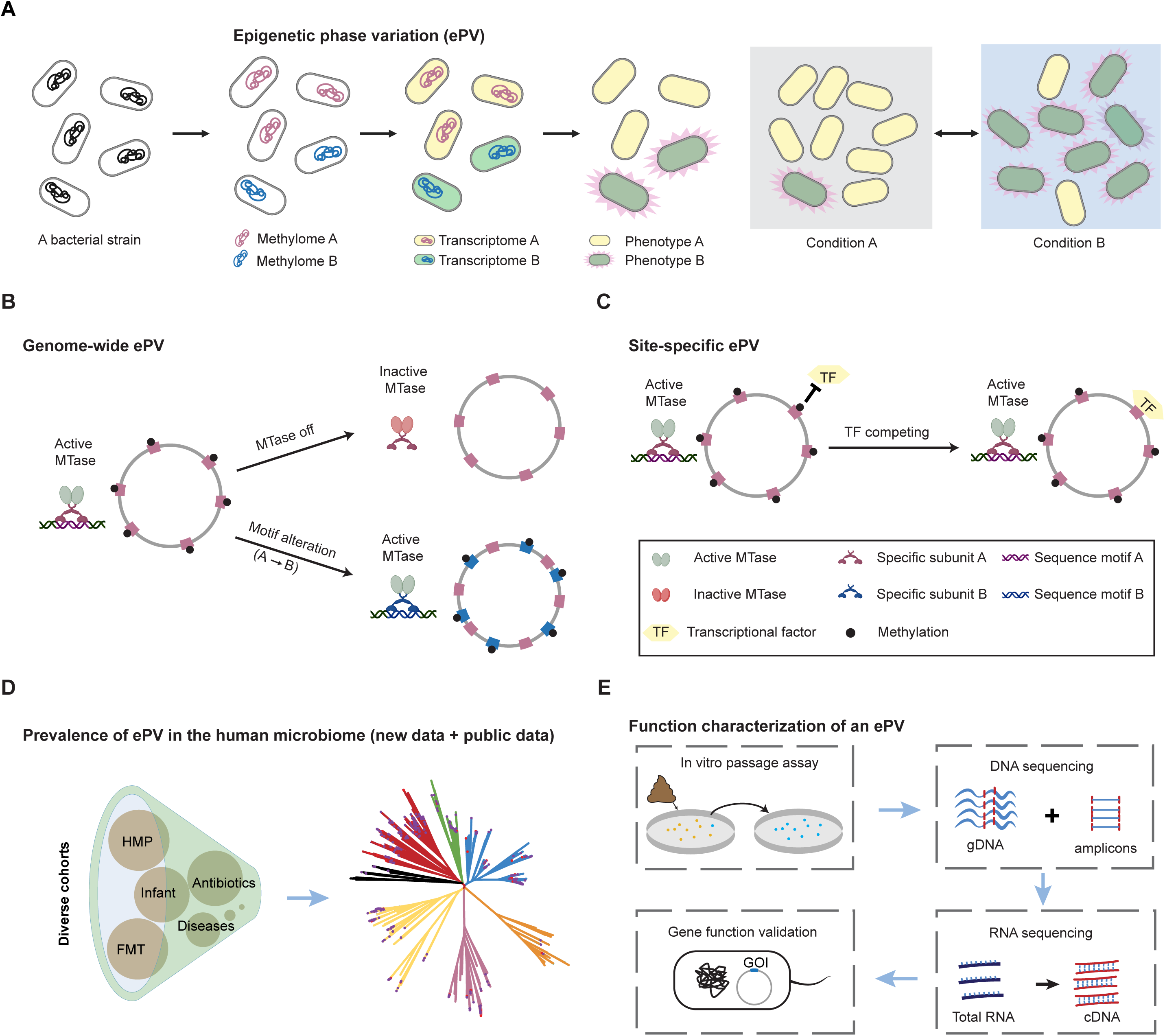
An integrated framework for epigenetic phase variation (ePV) detection and function validation from the human gut microbiome. (**A**) A specific bacterial strain may have sub-populations that exhibit heterogeneous methylation patterns, leading to distinct gene expression and phenotypes. This may enhance the ability of a single strain to adapt to different stress conditions. (**B**) Schematic diagram of genome-wide ePV events, mediated by the structural variation located within MTase genes (ON/OFF states) or their specificity units (alters recognition motifs). (**C**) Schematic diagram of site-specific ePV events, resulting from the competition between MTase and transcription factors (TFs). (**D**) Prevalence and phylogenetic distribution of ePV were explored across various healthy and clinical-related cohorts. Phylogenetic tree is color-coded at phylum level. (**E**) Workflow for functional analysis of genes involved in ePV events. Bacterial strains carrying the ePVs were isolated from fecal samples. An *in vitro* passage assay was performed under conditions mimicking the *in vivo* environment, followed by whole-genome shotgun and targeted DNA sequencing. MTase genotypes were quantified and associated with treatment conditions. RNA sequencing was conducted to profile transcriptomic differences between genotypes. Differentially expressed genes (DEGs) were overexpressed further in native strains or model species, and phenotypic tests were carried out to confirm whether the altered phenotype was due to the expression of the gene of interest (GOI).

ePVs characterized across diverse human pathogens can be categorized into two types: genome-wide or site-specific.^36–39^ A genome-wide ePV is usually mediated by genetic phase variation affecting an MTase or its corresponding specificity unit (which determines the recognition specificity, or sequence motif, of an MTase). In a diverse population, DNA MTase phase variation results in genome-wide activity of the MTase that alternates between ON/OFF states, leading to varying levels of methylation across the population, typically approaching either ∼100% or ∼0% methylation of its target motif in each bacterial cell (**Fig. 1B**).^33,38,39^ The phase variation of the specificity unit associated with a DNA MTase can create two or more alternative genome-wide recognition motifs (**Fig. 1B**).^30,40^ Both types of genome-wide ePVs can affect the expression of multiple downstream genes via epigenetic regulation, which provide greater flexibility than single-gene phase variation in creating transcriptional heterogeneity (e.g. *Streptococcus pneumoniae,*^41^ *Haemophilus influenzae,*^42^ *Helicobacter pylori,*^43^ *etc.*) and selective advantages to bacterial pathogens (e.g. *Mycobacterium tuberculosis*^44^). In contrast to genome-wide ePVs, site-specific ePVs involve a small number (one or few) methylation sites and are usually driven by the competitive binding between a DNA MTase and other DNA binding proteins such as a transcription factor (TF) (**Fig. 1C**). Two well-characterized examples are the ePV of the *pap* operon^45^ and the *agn43* gene^46^ in *Escherichia coli*, where the 6mA MTase targeting 5’-GATC-3’ sites compete with Lrp and OxyR, respectively. These site-specific epigenetic states are inheritable and reversible and have the advantage of mediating phase variation without altered genomic sequences, thereby mitigating mutation costs, for example, associated with antibiotic resistance.^47^ Both genome-wide and site-specific ePVs facilitate the manifestation of diverse phenotypic states within a single bacterial strain population, enhancing the ability to adapt to changing conditions.^48^

DNA methylation is highly abundant and diverse among the species constituting the human gut microbiome,^49^ yet the role of ePV and its impact on bacterial adaptation in the human gut remains largely unexplored. In this study, we aimed to address three major questions: (i) How prevalent are genome-wide and site-specific ePVs in the human gut microbiome? (ii) What are the functions of genes regulated by ePVs? (iii) How does an ePV enhance gut bacteria’s adaptation to environmental (e.g. antibiotics) and individual-specific (e.g. fecal microbiota transplantation, FMT) selective pressures?

To address these questions, we systematically investigated the occurrence and implications of ePV in the human infant and adult gut microbiome (**Fig. 1D-E**). Using long-read metagenomic and meta-epigenomic analysis, we detected genome-wide ePVs mediated by complex SVs in DNA MTases and their specificity units. We observed intra-strain ePVs associated with antibiotic use within individuals, as well as intra-strain ePVs between FMT donors and recipients. Furthermore, we analyzed a collection of publicly available short-read metagenomic sequencing datasets (2,345 unique samples) to estimate the prevalence of genome-wide ePVs mediated by structural variations in DNA MTases and their specificity units. In addition, we identified site-specific ePVs associated with probiotic engraftment and antibiotic treatment using single-molecule quantitative methylation analysis. Importantly, we observed that an *Akkermansia muciniphila* strain in an infant gut microbiome had an ePV detected following a course of amoxicillin (AMX); we recapitulated this shift *in vitro* and functionally characterized how this ePV enhances the adaptive capacity of the *A. muciniphila* strain in response to different antibiotics classes (**Fig. 1E**).

## Results

### Genome-wide epigenetic phase variations (ePVs) in the gut microbiome

To detect genome-wide ePVs, we performed joint metagenomic and meta-epigenomic analysis using long-read DNA sequencing (**Fig. S1A**; **Methods**) on both publicly available long-read metagenomic sequencing data and new data we generated for this study (**Fig. S1B**, **Table S1**). We focused on detecting structural variations (SVs) involving DNA MTases and/or their corresponding specificity units, including invertible repeats, and complex SVs spanning one or multiple coding regions (**Methods**). These SVs have been linked to genome-wide ePV in bacterial pathogens^41–44^ but not in the gut microbiome. While short-read DNA sequencing can detect invertible repeats, long-read DNA sequencing can resolve complex SVs by assigning individual long reads to a different possible genome rearrangement built from each reference genome (**Fig. S1B**, **Methods**).^22,50–52^ We used long-read mapping tools (pbalign / NGMLR^52^) with additional quality controls to ensure read homogeneity at genomic regions flanking the SVs that reflect intra-strain variation (**Methods**). For example, from the microbiome sample from a healthy adult,^53^ we detected a complex SV within an MTase in *Phocaeicola vulgatus* (formerly *Bacteroides vulgatus*), mediated by three paired/matching inverted repeats, resulting in three possible genotypes (g1-g3; **Fig. 2A**). Individual long reads can be unambiguously assigned to each of the three genotypes, facilitating genotype-specific quantitation (**Fig. 2A**). Across the 221 long-read metagenomic samples analyzed, we detected 294 SVs involving DNA MTase and/or specificity units (**Table S2**), which encode 2-8 alternative genotypes based on long-read mapping analysis (**Fig. 2B**, **Fig. S1C**). One of these SVs was detected from *Gemmiger qucibialis* [within class Clostridia] in the context of an FMT trio (donor: D1; donor 5 years: D1.5y; and recipient post-FMT: R1-postFMT; **Fig. 2C**). Specifically, this SV has two alternative genotypes (g1 and g2) which demonstrate different patterns when colonizing donor and recipient. While g1 is the dominant genotype in the *G. qucibialis* strain in the FMT donor sample (very stable over a five-year time window, **Table S3**), g2 becomes the dominant genotype in the recipient one year after FMT (**Fig. 2C**, **Methods**), suggesting differing host selective pressures between the FMT donor and recipient. We built on our recently described method for precise tracking of individual strains, performed rigorous long-read mapping analysis and strain-level *de novo* metagenomic assembly (**Fig. S2**, **Supplementary Data**) to confirm the SV in *G. qucibialis* was indeed from the same donor strain that stably engrafted the recipient post-FMT.^54^

**Figure 2.**
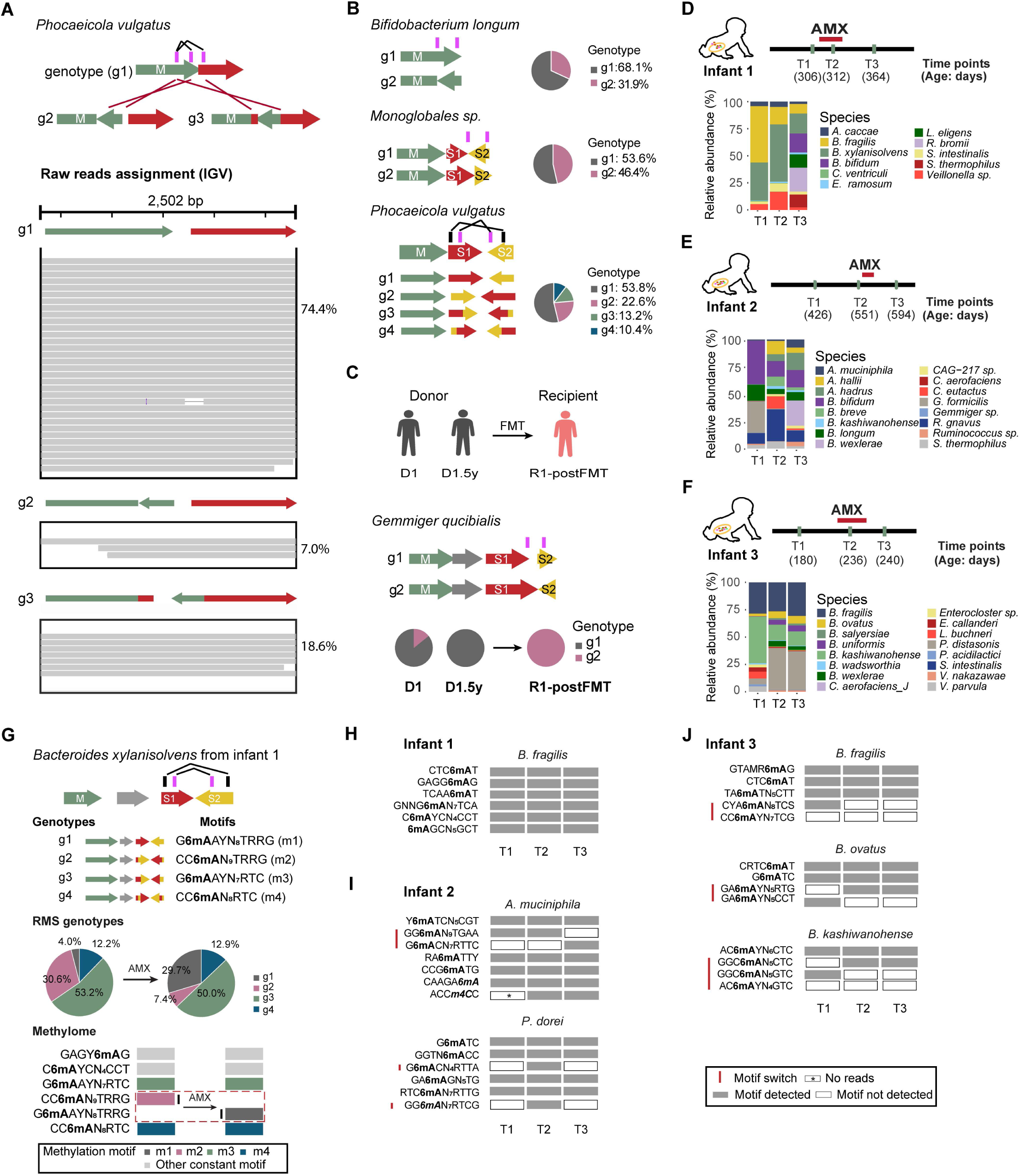
Long-read metagenomic sequencing deciphers complex ePV events. (**A**) An ePV observed in a *Phocaeicola vulgatus* strain from a healthy adult gut microbiome, mediated by three paired/matching inverted repeats (inverted repeats), resulting in three distinct MTase genotypes (g1-g3). The alignment plots show raw reads aligned with their corresponding reference genomes. The proportion of each genotype is displayed on the right side of the read alignment chart. (**B**) Additional ePV events were identified in various bacterial species. Gene diagrams illustrate the rearrangement of MTase gene fragments, while pie charts show the proportion of each genotype variant. (**C**) ePV events identified in an FMT cohort. The top panel presents metadata explaining sample and subject information (see **Table S3**). The bottom panel shows an ePV in a *Gemmiger qucibialis* strain, detected in both donor and post-FMT samples. (**D-F**) Metagenomic analysis of fecal samples from three infants in the ECAM cohort. The top panel presents the timeline of sample collection and amoxicillin (AMX) exposure. The bottom panel displays species-level relative abundance profiles, including only species with a sequencing depth > 10X. (**G**) An ePV event was observed in a *Bacteroides xylanisolvens* strain, mediated by two pairs of IR features, resulting in four specificity unit genotypes (g1-g4). The genotype was profiled in pre- and post-AMX treatment samples, alongside their corresponding methylome (m1-m4). AMX treatment suppressed the population of g2 (highlighted in purple) while promoting the growth of g1 (highlighted in grey). Simultaneously, the methylome shifted from m2 to m1, mirroring the genotype changes. (**H-J**) Additional methylation changes and ePV events in different bacterial species in Infant 1 (**H**), Infant 2 (**I**) and Infant 3 (**J**). The methylation profiles were generated based on the single-site methylation analysis (see **Methods**) and manually curated. (**A-C**, **G**) M: MTase represented by green arrows; S1, specificity unit 1 represented by red arrows; S2, specificity unit 2 represented by yellow arrows. The pink and black rectangles above the gene schematic diagram are inverted repeats.

Because PacBio sequencing data support direct detection of 6mA and 4mC, *de novo* methylation motif and quantitative methylation analysis at single molecule resolution (**Methods**),^29,55,56^ we were able to directly assess ePVs mediated by the SVs of DNA MTase genes and/or their specificity units. We focused on nine fecal samples (from three infants, ages 1-3) collected as part of the Early Childhood Antibiotics and the Microbiome (ECAM) study.^3^ The three infants were selected based on their exposure to amoxicillin treatment, fecal samples availability both before and after administration of the antibiotics, which had substantial impact on microbiome composition in each case (**Fig. 2D-2F**). To illustrate how the SV of an MTase and specificity unit mediated a genome-wide ePV, we describe one example detected from a *Bacteroides xylanisolvens* strain in Infant 1. Two pairs of inverted repeats in the specificity unit create four possible genotypes via genome rearrangement, with four expected methylation recognition motifs (**Fig. 2G**). For these four genotypes, *de novo* methylation analysis resolved specific methylation motifs, and ePVs estimated from methylation analysis correlate with the genotype frequencies estimated from long-read mapping (**Fig. 2G**; **Methods**). Across the three infants, we observed bacterial strains whose methylation motifs were stable over time (**Fig. 2H-2J**), as well as those with genome-wide ePV in at least one methylation motif following exposure to amoxicillin (**Fig. 2H-2J**), which suggests possible antibiotics-induced selective pressure of genome-wide ePVs.

Although long-read sequencing is powerful for metagenomic SV analysis, most existing microbiome studies were based on short-read sequencing platforms. While short-read sequencing has limited ability to resolve complex SVs, they are reliable for detecting relatively simple SVs (e.g. with only two alternative genotypes).^21,22^ Building on PhaseFinder,^21^ we adapted a pipeline for detecting putative ePVs from short-read metagenomic shotgun sequencing data (**Fig. 3A-3B; Methods**) and applied it to 2,345 unique metagenomic samples from nine studies including the Human Microbiome Project^57^ and additional cohorts (**Table S4**). Across 1,308 unique species, we detected 2,556 putative genome-wide ePVs with at least one alternative genotype based on SVs in MTases or specificity units (**Fig. 3C**). Four orders (Clostridiales, Bacteroidales, Enterobacterales, and Bifidobacteriales) were enriched for ePVs (**Fig. 3C**; **Fig. S3A-S3B**), although we observed broad prevalence of potential ePVs (inverted repeats in MTase or specificity units, with neighboring invertase) across many other orders (**Fig. 3C**). Among the detected putative ePVs, some are associated with the use of antibiotics. For example, a *B. uniformis* strain from an infant cohort^58^ has a putative ePV with two alternative MTase genotypes, which had clear changes upon amoxicillin treatment, across four infants (**Fig. 3A, 3D**). The MTase gene and SV involved in this ePV was detected in 23 of 153 complete *B. uniforms* genomes available at NCBI, with each of these 23 strains exhibiting one of the two ePV genotypes (**Fig. 3E**).

**Figure 3.**
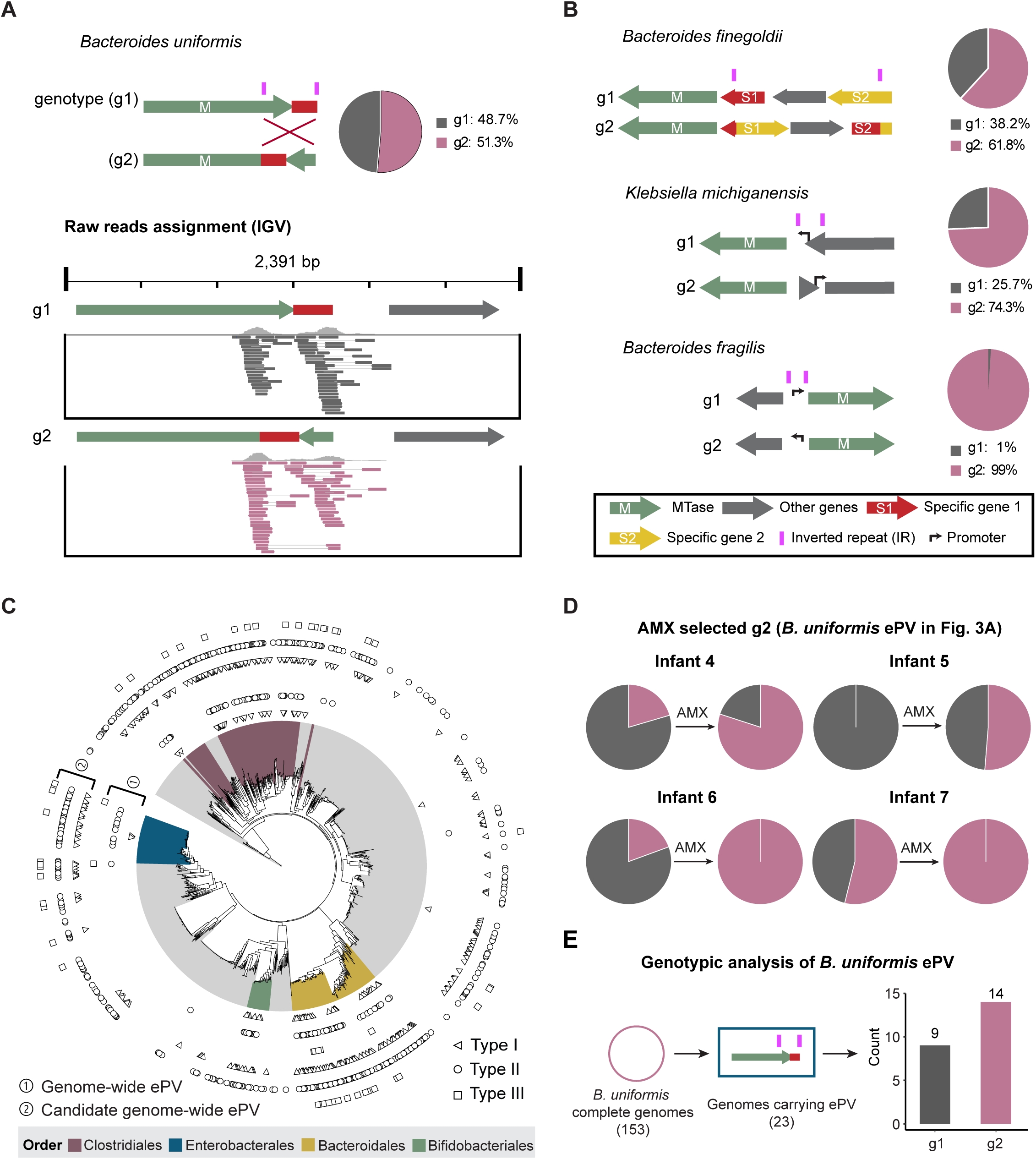
Genome-wide ePV events are widespread across different cohorts and associated with clinical treatments. (**A**) An ePV observed in a *Bacteroides uniforms* strain from an adult gut microbiome. Due to read length limitation, short-read data can only detect simple ePV (or “inverton”, as defined in a previous study^21^), which are mediated by a pair of IR features and result in two distinct MTase genotypes (g1, g2). The proportion of each genotype is shown to the right of the gene diagram. The read alignment chart illustrates how paired-end (PE) reads were used to detect the inverton-like ePV. (**B**) Additional inverton-like ePVs detected across various bacterial species. Gene diagrams show the rearrangement of MTase gene fragments, while pie charts depict the proportion of each genotype variant. (**C**) Phylogenetic tree of inverton-like ePVs. Potential ePV (based solely on the reference genome) and confirmed ePV (supported by both the reference genome and metagenomic read) across three types of Restriction-Modification Systems (I, II, III) (**Fig. S3**) and are projected onto the phylogenetic tree. ePV enriched orders are highlighted according to the colors scheme in the legend. (**D**) Response of *B. uniforms* ePV to AMX treatment. The *B. uniformis* ePV mentioned in Fig. 3A was detected in four infants from the DIABIMMUNE cohort, both pre- and post-AMX treatment. The MTase genotype shifted from g1 to g2 in response to treatment. (**E**) Prevalence analysis of the *B. uniformis* ePV. Among 153 complete *B. uniforms* genomes from NCBI, 23 were found to carry this ePV. The bar plot on the right shows the frequency of the dominant ePV genotype across the 23 genomes. (**A**, **B**, **E**) M: MTase represented by green arrows; S1, specificity unit 1 represented by red arrows; S2, specificity unit 2 represented by yellow arrows. The pink rectangles above the gene schematic are inverted repeats.

### Site-specific epigenetic phase variations (ePV) in the gut microbiome

Differing from the genome-wide ePVs resulting from changes in MTase activity or specificity, site-specific ePVs involve only one or several methylation sites, and are usually driven by the site-specific binding competition between a DNA MTase and another DNA binding protein such as a TF (**Fig. 4A**). For a specific sequence motif recognized by a DNA MTase in a bacterial strain, while most motif sites are stably methylated (∼100%) across bacterial cells at a certain condition, a few motif sites are only methylated in a small percentage (for example,

**Figure 4.**
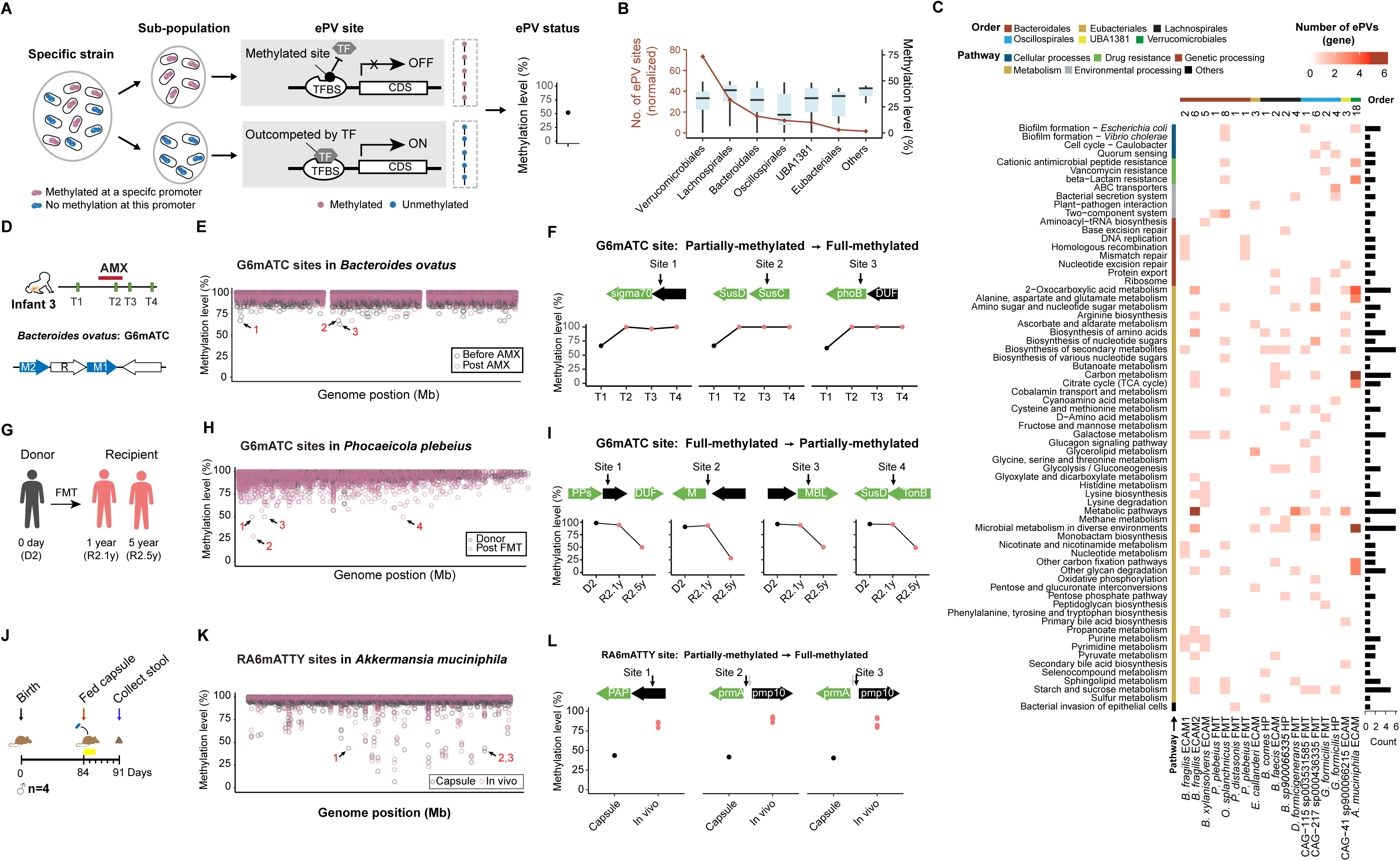
Site-specific ePV events are widespread across different cohorts and associated with bacterial adaptation to environmental stress. (**A**) For a specific bacterial strain, only a subset of cells exhibit methylation at a specific promoter, while the remainder remain unmethylated (left panel). The schematic in the middle panel depicts a transcription factor (TF) competing with an MTase at this specific site. The ePV site (highlighted within the dashed rectangle) was identified from multiple molecules using single-molecule methylation analysis. The right panel illustrates the deconvolution results obtained from this single-site methylation analysis. (**B**) Summary of all ePV sites detected in this study. Values were normalized based on the strain number within the order (diamond points highlighted in red). The methylation percentage within the order were also determined (box plot). Orders with fewer than five strains were merged into the ’Others’ category. (**C**) Function categories of the genes affected by ePV sites. Heatmap showing the number of genes regulated by ePV, with host strain taxonomy (strain) as column label and gene function category (pathway) as row label. The color scale represents the number of genes, indicating the occurrence of each functional category per strain. (**D**) The top panel presents the time-line of AMX administration and sample collection for Infant 3. The bottom panel shows the involved methylation-related genes in a *B. ovatus* strain from Infant 3. (**E**) Distribution of partially methylated G6mATC sites across the *B. ovatus* genome. The genome consists of three contigs, randomly ordered along the x axis. The top three changed partially methylated sites were selected and labeled with numbers. (**F**) Changes in partially methylated G6mATC sites in response to AMX treatment. The motif sites were examined on both strands, and only sites that showed partial methylation on both strands were considered final ePV sites. The same criteria were applied to all other partially methylated sites in Fig. 4I,**L**. The gene diagram shows the location of each site within the affected gene. **(E,F)** Pre-(black) and post-AMX (red) methylations were highlighted. (**G**) Sample collection information for Donor 2 and Recipient 2 from FMT cohort. (**H**) Distribution of partially methylated G6mATC sites across the *Phocaeicola plebeius* genome. The top four partially methylated sites were selected and labeled with numbers. (**I**) Changes in partially methylated G6mATC sites in response to FMT. The gene diagram shows the location of each site within the affected gene. **(H**,**I)** Pre-(black) and post-FMT (red) methylations were highlighted. (**J**) Design and sample collection information for Akk-fed mouse assay. (**K**) Distribution of partial methylated RA6mATTY sites across the *A. muciniphila* genome. Three of the partially methylated sites, specifically lowly methylated in capsule, were identified and labeled with numbers. (**L**) Changes in partially methylated RA6mATTY sites from capsule to *in vivo* colonization (mouse stool). The gene diagram shows the location of each site within the affected gene. **(K**,**L)** Capsule (black) and *in vivo* (red) methylations were highlighted.

<50%) of bacterial cells (**Fig 4A**). We built on previous work for PacBio DNA sequencing-based single-molecule methylation analysis,^29,59–61^ now with enhanced quality control to ensure site-specific ePV reflects variation within a single strain rather than a mixture of different strains (**Methods**). Methods evaluation supported that the pipeline can reliably quantify the extent of partial methylation of individual methylation motif sites (**Fig. S4A**).

We performed quantitative site-specific ePV detection on 12 ECAM infant samples and **8** FMT fecal samples, and discovered 885 site-specific ePVs with < 50% site-specific methylation across 32 strains (representing 27 unique species), including some motif sites that were mostly (<25% or even <10%) unmethylated (boxplot in **Fig. 4B**; **Fig. S4B**). The ePV sites were classified as either intergenic or intra-genic based on their positions relative to protein-coding genes. Of the 885 site-specific ePVs, 174 (19.7%) are in intergenic regions (**Fig. S4C**). This enrichment of ePV sites in intergenic regions (Chi-square *p* < 10e-5) is consistent with the proposed mechanism of site-specific competitive binding between MTases and transcription factors (or repressors),^45,46^ although intragenic DNA methylation might also be associated with the regulation of gene expression in bacteria.^29,31^ The genes flanking site-specific ePVs have functions across diverse categories related to metabolic pathways, cellular processes, and environmental processing including several genes associated with host interactions, antibiotic resistance, and biofilm formation (**Fig. 4C**).

To determine whether certain site-specific ePVs are associated with the selective pressure of environmental factors, we examined the 12 ECAM cohort fecal samples from infants with amoxicillin exposures as described above. We detected ePVs with >40% differences in their methylation status between two consecutive sampling time points. In Infant 3 (**Fig. 4D**), we found three site-specific ePVs in *Bacteroides ovatus* (**Fig. S4D**) whose methylation levels significantly increased upon the infant’s treatment with amoxicillin (**Fig. 4E-4F**). The first ePV is in the 5’ upstream of the essential sigma factor σ^70^, which regulates genes involved in basic cellular processes including protein synthesis, energy production, and cell wall biosynthesis.^62^ The second ePV is located near the genes encoding SusC/SusD family outer membrane proteins, which have been identified in various *Bacteroides* species as phase-variable factors.^63,64^ The third ePV is located in the *phoB* gene, which encodes a response regulator protein essential for the PhoBR two-component signal transduction system. This system regulates the expression of multiple genes under phosphate-limiting conditions^65^ and plays a critical role in biofilm formation and environmental stress response.^66^ The presence of site-specific ePVs in these genes likely contributes to transcriptional heterogeneity, providing additional adaptive capacity under amoxicillin exposure. Similarly, we examined the FMT fecal samples as described above. Between the FMT donor and post-FMT recipient samples **(Fig. 4G)**, we detected ePV sites with differentially methylation ≥ 50% (**Fig. 4H**). In donor 2, we found four site-specific ePVs in *Phocaeicola plebeius* whose methylation levels had major decreases 5 years after FMT (**Fig. 4H-4I**, **Fig. S4E**). One of the four sites is located in the *MBL* gene encoding a β-lactamase, which are emerging as the most notable resistance determinants in Enterobacteriaceae^67,68^, while another site is located between *susD* and *TonB*, both were reported to be phase variable sites in several *Bacteroides* species.^54,63,64^

To determine whether site-specific ePVs may be affected by selective pressure in the gut environment differing from *in vitro* culture, we fed a commercially available probiotic capsule [Glucose Control, Pendulum] with five strains (*A. muciniphila*, *Bifidobacterium longum*, *Clostridium beijerinckii*, *Clostridium butyricum*, and *Anaerobutyricum hallii*) to Germ-free B6 mice and collected fecal samples at day 7 (**Fig. 4J**). PacBio long-read metagenomic sequencing data showed that *A. muciniphila* stably colonized all four tested mice (**Fig. S5A**). Site-specific methylation analysis found 38 ePVs across 15 unique RA6mATTY sites in the probiotic *A. muciniphila* strain, showing >50% differential methylation between the inoculum (capsule) and the fecal (*in vivo*) samples (**Fig. 4J-4L**). These findings were further supported by independent validation based on restriction digestion (**Fig. S5B**). In contrast, no ePV events were observed at A6mACN_6_TAC sites (**Fig. S5C**). Five of the 16 genes flanking the 15 unique RA6mATTY sites exhibited host colonization defects: *hxlR*, *MASP*, *GLUL*, *fhaB* and Amuc_0952.^69^ Additionally, another gene identified exclusively in this study, *pmp10*, is also associated with bacteria-host interaction^70,71^ (**Fig 4L**). Some other genes involved in these site-specific ePVs are known to be associated with stress responses that help bacteria adapt to challenging conditions (*prmA*^72^*, PAP*^73^, **Fig. 4L**). The selection for *A. muciniphila* cells with methylated ePV states of these genes suggest they are associated with the engraftment of the probiotic strain in the gut environment.

### Genes, functions and selective advantage mediated by a genome-wide ePV in an *A. muciniphila* strain

Next, we focused on *A. muciniphila,* a promising next-generation probiotic associated with several positive health outcomes and drug responses.^74–78^ This species was selected for its prevalent genome-wide and site-specific ePVs (**Fig. 2I**, **4K-4L**) and its response to amoxicillin exposure observed in an infant fecal sample (**Fig. 5A**). This strain encodes a MTase specificity unit with three nested inverted repeats, creating eight different genotypes (g1-g8) and six unique methylation sequence recognition motifs, corresponding to six epigenetic phases m1–m6 (see **Fig. 5A; Methods**). Upon the use of amoxicillin, the subpopulation of g2 increased from 11.1% to 26.5% (**Fig. 5A; Methods**), consistent with a genome-wide loss of 6mA at m1:GG6mAN^9^TGAA sites and genome-wide gain of 6mA at m2:G6mACN_7_RTTC sites (**Fig. S6A**) due to the genome-wide change in MTase sequence recognition specificity.

**Figure 5.**
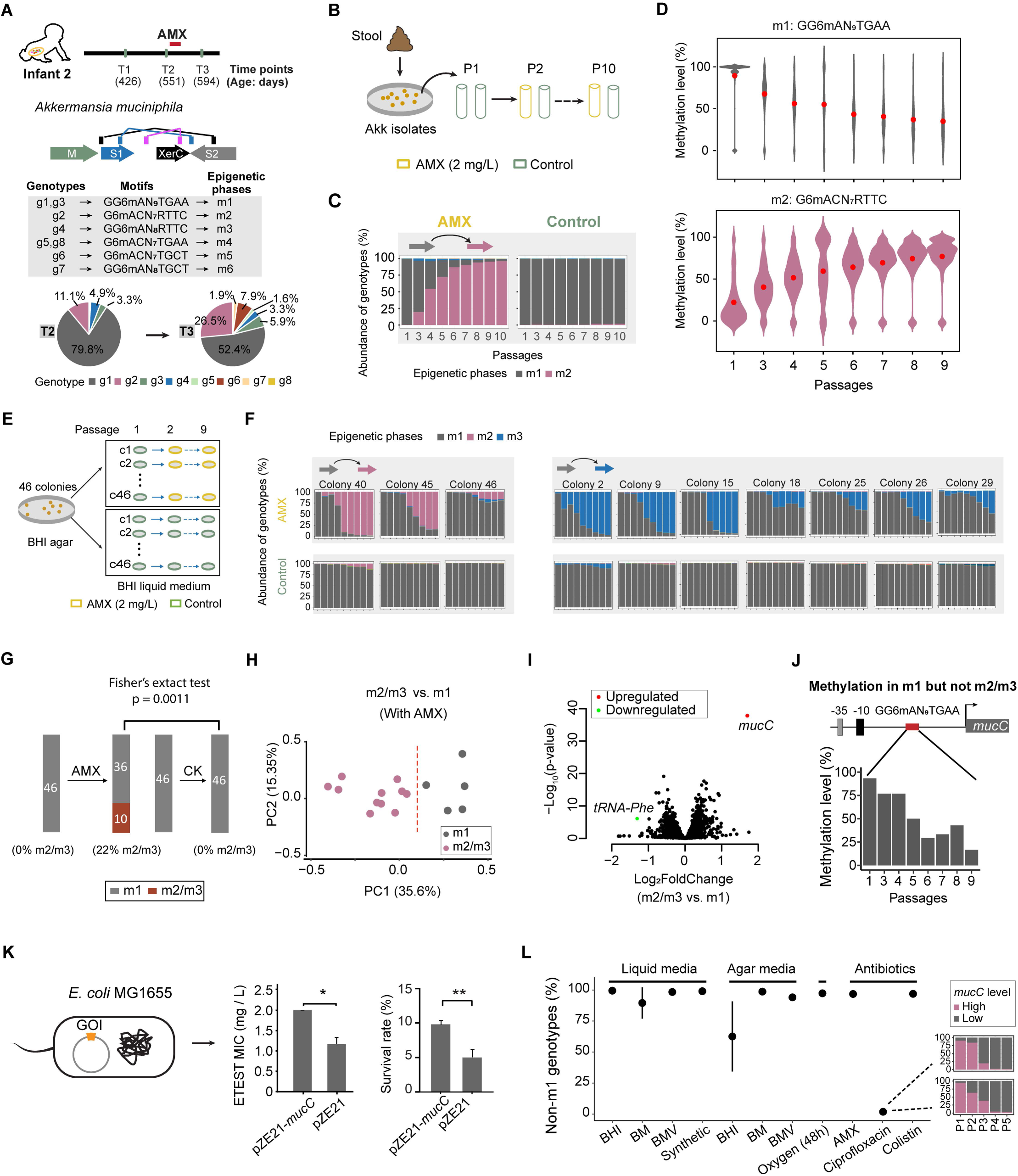
Characterization of a complex genome-wide ePV and its role in *Akkermansia* tolerance to AMX. (**A**) An ePV event observed in an *A. muciniphila* strain from Infant 2 in the ECAM cohort, mediated by three pairs of IR features, resulting in eight MTase genotypes (g1-g8) and six unique methylation sequence recognition motifs. The genotype was profiled in pre- and post-AMX treatment samples, AMX treatment suppressed the population of g1 (highlighted in grey) while promoting the growth of g2 (highlighted in purple) or other non g1 populations. (**B**) Experimental design of the *in vitro* passage assay. *A. muciniphila* was isolated from the fecal sample of Infant 2 (hereafter referred to as Akk_ECAM). After recovery on the BHI agar plates, a culture derived from a single colony was subjected to 2 mg / L AMX treatment for 10 passages. Cultures without AMX served as negative controls. (**C**) MTase genotype profiling using targeted sequencing. In response to AMX treatment, the MTase genotype changed from g1 dominance to g2 dominance, whereas g1 remained dominant through passage 10 in the absence of AMX. (**D**) Methylation motif changed in the Akk_ECAM strain upon AMX treatment. Genomic DNA was extracted, and whole genome shotgun sequencing was performed using Pacbio Sequel II platform. Single-molecule methylation analysis was then conducted to investigate the methylation level of the two motifs of interest. The results align with the *in vivo* findings, where AMX treatment decreased in the g1 methylation subpopulation (highly methylated at GG6mAN_9_TGAA) and increased the non-g1 methylation subpopulation (highly methylated at G6mACN_7_RTTC). (**E**) Experimental design of the 2^nd^ batch of the *in vitro* passage assay. After recovery on BHI agar plates, 46 colonies were subjected to 2 mg / L AMX treatment for nine passages. Culture without AMX served as negative controls. (**F**) MTase genotype profiling with targeted sequencing. The MTase genotype changed from g1 dominance to non-g1 (g2-g4) dominance in response to AMX treatment, while g1 remained dominant through passage 9 in the absence of AMX treatment. (**G**) Summary of the 2^nd^ batch *in vitro* passage assay. Genotypes were determined by targeted sequencing and categorized into two groups, g1 and non-g1 (g2-g4). Fisher’s exact test was used to assess the statistical significance of the difference in proportions between pre- and post-AMX treatment. The results indicate a significant association (p = 0.0037), suggesting that g2-g4 genotypes were selected by AMX treatment. (**H**) Principal Component Analysis (PCA) analysis of RNA-seq-based expression data from in vitro passage samples. Under AMX treatment, g2/g4 samples were clearly separated from g1/g3 samples. Overall sample variation was evaluated using PCA, with the explained variation labeled as percentages on the axes. (**I**) DEGs were identified by comparing g1/g3 and g2/g4 samples under AMX treatment. Thresholds for DEGs were set at log_2_ fold changes > 1 or < -1, with an adjusted p value < 0.05. *mucC* was the only upregulated gene (highlighted in red). *tRNA-Phe* was the only downregulated gene (highlighted in green). (**J**) The promoter elements and m1 methylation motif (GG6mAN_9_TGAA) were represented in the diagram. The fraction of m1 methylation motif in the promoter of *mucC* decreased with AMX treatment. (**K**) AMX tolerance of *E. coli* and overexpression of the *mucC* gene. The *mucC* gene was codon-optimized, synthesized and fused with a 6 × HIS tag at the N-terminus. It was inserted into a medium-copy plasmid with a constitutive promoter, and AMX tolerance was assessed using the ETEST and CFU counting methods respectively. Each strain was tested in triplicate. (**L**) MTase specificity reversibility assay. g2/g4 dominant strains from the 2^nd^ batch of the *in vitro* assay were treated under various conditions. The point plot shows the targeted sequencing results, while the zoomed-in bar plot indicates that ciprofloxacin treatment can reverse non-g1 (g2/g4) genotypes back to g1 genotype. Each condition was tested in duplicate.

To characterize the impact of this genome-wide ePV on bacterial fitness, we first isolated the *A. muciniphila* strain (**Methods**) and selected a clone (hereafter referred to as Akk_ECAM) that is dominated by the m1 genotype representing the same ePV state as in the infant’s fecal sample before amoxicillin exposure (**Fig. S6B**). After ten passages of sub-cultures in liquid media with amoxicillin (**Fig. 5B**; **Fig. S6C**; **Methods**), we observed gradual population-level change from ePV phase m1 to m2 (**Fig. 5C**) along with genome-wide methylation changes from GG6mAN_9_TGAA to G6mACN_7_RTTC (**Fig. 5D**). These gradual epigenomic changes were not observed in controls without amoxicillin (**Fig. 5B-D**). The directionality of the changes (the m2 ePV state selected by amoxicillin) is consistent with the observations from the infant’s fecal samples (**Fig. 5A**). However, in two additional clones, we did not observe the same changes in genome-wide ePV, suggesting that *A. muciniphila* may adapt to amoxicillin through multiple mechanisms.^79^ Therefore, we performed a more comprehensive characterization of this ePV across a larger number (n=46) of m1 clones (**Fig. 5E**). Consistent with our initial observations, there was a variability among these 46 clones in terms of how the ePV state changed during the nine passages in the presence of amoxicillin: three clones changed to m2 and seven changed to m3 (**Fig. 5F**); the remaining clones remained as m1. In contrast, in the absence of amoxicillin, all clones remained at m1 after the passages. With the initial abundance of the non-m1 ePV state < 0.1% at the beginning of the passage, that 10 out of 46 clones changed their ePV states in the presence of amoxicillin is statistically significant (Fisher’s exact test *p* = 0.0011; **Fig. 5G**), suggesting that this genome-wide ePV may serve as one of several mechanisms that enhance the tolerance of the *A. muciniphila* strain to amoxicillin. Because m1 preserves methylation motif GG6mAN_9_TGAA while m2 and m3 both lose the methylation motif (**Fig. 5A**), we hypothesized that the loss of GG6mAN_9_TGAA methylation in m2 and m3 influences the expression of certain *A. muciniphila* genes that increases its tolerance to amoxicillin.

To test this hypothesis, we selected nine clones in total with preserved (m1) or lost (m2, m3) GG6mAN_9_TGAA methylation and used RNA-seq to profile their gene expression in the last two passages in the presence of amoxicillin (**Fig. S7A**). The RNA-seq data analysis showed that the global gene expression of the m1 clones and m2/m3 clones have subtle but consistent differences (**Fig. 5H**). Differential gene expression analysis identified a single gene [Amuc_0914 homolog, herein named *mucC*, for *muciniphila* gene from cupin family] that is significantly (negative binomial test, *p* = 3.76E-07) and consistently up-regulated when GG6mAN_9_TGAA methylation is lost in m2 and m3 clones (**Fig. 5I**, **Fig. S7B**). Consistently, the promoter of *mucC* contains a GG6mAN_9_TGAA methylation site between its -35 region and translation initiation site (**Fig. 5J**) suggesting methylation-mediated regulation of gene expression. To test if some genetic variations might be selected by amoxicillin, we also performed whole genome shotgun sequencing on m2/m3 colonies: although we found several genetic variations across the clones, none were associated with the amoxicillin treatment (**Fig. S7C**). These findings suggest that the loss of GG6mAN_9_TGAA methylation in the promoter of *mucC* gene is mediated by the genome-wide ePV changing from m1 to m2/m3, upregulating *mucC* expression, which enhances the tolerance of this *A. muciniphila* to amoxicillin.

The *mucC* gene is highly conserved across *A. muciniphila* strains (**Fig. S8A**, **Table S5**) encoding a small protein characterized by a cupin fold,^80^ which consists of a conserved β-barrel structure (AlphaFold prediction,^81,82^ **Fig. S8B**). MucC homologs are also found in other bacterial species (**Fig. S8C**). In *A. muciniphila*, *mucC* is immediately adjacent to Ribonuclease R (rnr), a member of the RNase II superfamily (**Fig. S8D**) that hydrolyze RNA in the 3’ - 5’ direction. RNase R is associated with selective mRNA degradation, particularly of mRNAs that lack a stop codon in bacteria^83^ and are expressed under stress conditions.^84^ Because the genetic tools of *A. muciniphila* are still limited^69^, we expressed *mucC in trans* in *E. coli* cells (pZE21-*mucC*) and confirmed its protein expression by mass spectrometry (**Methods**; **Fig. S9A-S9D**). By comparing *mucC*^+^ *E. coli* (pZE21-*mucC*) with negative controls (empty plasmid, pZE21), *mucC* expression conferred significantly greater tolerance to amoxicillin based on their MICs and survival rate (**Fig.5K**).

Finally, because ePV enhances non-genetic clonal heterogeneity and empowers the adaptive capacity of single cells in a population,^39,41–44^ we hypothesized that while the ePV state with high *mucC* expression enhances tolerance to amoxicillin, other conditions may favor an ePV state with low *mucC* expression. High basal level of *mucC* expression doesn’t create a disadvantageous fitness cost *in vitro*, as we found that the removal of amoxicillin didn’t select for the m1 ePV state with lower *mucC* expression (**Fig. S10**). However, after testing several additional stress conditions, we discovered that ciprofloxacin selects the m1 ePV state with lower *mucC* expression (**Fig. 5L**, **Table S7**), demonstrating a bet hedging strategy of dynamic adaptation to different stresses. Further examination led to the observation that the selection of m1 epigenetic phase (low *mucC* expression) by ciprofloxacin varied between clones and replicates (**Fig. S11A**), which can be partially explained by additional SNPs (**Fig. S11B**) or structural variations (**Fig. S11C**). Ciprofloxacin is a fluoroquinolone antibiotic that targets bacterial DNA gyrase and topoisomerase IV, leading to double-stranded DNA breaks. This DNA damage can trigger the bacterial SOS response, resulting in increased mutation rates and the potential emergence of resistance-conferring mutations. Nevertheless, our findings suggest that ePV provides an alternative pathway for the *A. muciniphila* strain to tolerate ciprofloxacin (**Fig. S11A**). These observations suggest that the *A. muciniphila* strain exhibits clonal heterogeneity involving both genetic and epigenetic variations, which collectively enhance its adaptation to antibiotics.

Overall, our findings indicate that a complex ePV detected in an *A. muciniphila* strain that originated from an infant gut microbiome enhances clonal heterogeneity via regulating *mucC* gene expression with several ePV states associated with high *mucC* expression selected by amoxicillin and low *mucC* expression selected by ciprofloxacin. The reversible selection by two different antibiotics highlights the advantage of ePV to empower the same *A. muciniphila* strain to epigenetically adapt to different stress conditions. Noteworthy, this evolutionary strategy, where a small subpopulation generates ePV heterogeneity sufficient to increase population-level adaptation across various stress conditions, underscores the adaptive advantage at the population level rather than individual cells.

## Discussion

Our study highlights the prevalence of ePV across diverse bacterial species within the gut microbiota, underscoring the role of DNA methylation in enhancing clonal heterogeneity and enabling strains to adapt to environmental stressors such as antibiotic treatments or when colonizing new hosts after FMT.

We leveraged long-read metagenomic data to simultaneously detect complex SVs and DNA methylation, providing a detailed view of the epigenetic landscape of bacteria within gastrointestinal tract. Furthermore, our analysis of a collection of short-read metagenomic data has highlighted SVs near MTase and specificity units, suggesting the broad presence of ePV in various human cohorts. As long-read sequencing is more increasingly used in microbiome studies, we anticipate that our findings will encourage a new dimension to characterize how gut bacteria adapt to various stress conditions and host interactions through epigenetic mechanisms. The genes associated with ePVs in our analyses suggest that the biological functions mediated by ePV are broad, impacting microbe-host interactions, antibiotic resistance, and the metabolic capacities of gut bacteria. Importantly, the gene functions implicated by ePVs require further investigation to elucidate their mechanistic contributions. In pursuit of this, we have conducted a detailed characterization of an ePV identified in an *Akkermansia muciniphila* strain, revealing a specific gene that is regulated by DNA methylation and enhances tolerance to amoxicillin. As genetic tools for *Akkermansia muciniphila* continue to evolve, future research will likely uncover additional mechanistic insights into how this and other ePVs detected play a role in the adaptability and functionality of this vital commensal species.

Compared to genetic phase variation, ePV offers several advantages in enhancing clonal heterogeneity. The reversibility of ePV, without altering DNA sequence or incurring mutation costs serves as an additional way for individual bacterial strains to adapt to diverse stresses. Furthermore, genome-wide ePV can simultaneously regulate multiple genes, which provides coordinated activation or inhibition of multiple functionally related genes.^30,33^ As shown in bacterial pathogens, regulation by global epigenetic modification highlights an intermediate state toward long-term adaptation to stress conditions and interactions with the human immune system.^26,79,85^ The diverse functions of the genes associated with both genome-wide and site-specific ePVs suggest that ePVs may regulate critical processes of gut bacteria to enhance their adaptation and resilience in the changing environment.

The ePV characterized in the *A. muciniphila* strain that regulated differential expression of the *mucC* gene upon exposure to two common antibiotics, amoxicillin and ciprofloxacin **(Fig. 5)**, is a great example of a new epigenetic bet-hedging mechanism of commensal bacteria. Our results indicate that in the human gut, ePVs may help bacterial populations regain heterogeneity after bottlenecks encountered during colonization of a new host or severe perturbations due to antibiotic exposures. These ePV-driven regulatory mechanisms open new opportunities for targeted epigenetic interventions to improve desired functions of beneficial bacteria. For example, by manipulating ePV, we may strategically boost the resilience and functional capabilities of beneficial bacteria, which might improve the success rates of probiotic engraftment and the efficacy of treatments for microbiota-associated conditions.

Despite that in our study, the epigenetically regulated *mucC* expression was an important factor enhancing tolerance of the *A. muciniphila* strain to amoxicillin, we observe certain variations across all clones, which suggest that the epigenetic bet hedging is not the only mechanism for the *A. muciniphila* strain to adapt to antibiotic or other stressors. As reported in other studies, genetic mutations can also confer increased fitness in bacterial species. For instance, a subpopulation shift in specific clones has been reported in *B. thetaiotaomicron*, which was associated with resistance to oxidative stress through alterations in the sequence of the IctA protein.^86^ Our finding highlights that ePV is one additional way to study clonal heterogeneity and should be considered within a systemic view that includes other genetic, transcriptional, and translational forms of regulations.^15,21,22,57^

## Data availability

All sequencing data generated in this study have been submitted to NCBI SRA with the accession number PRJNA1207694, and will be made publicly available upon publication.

## Acknowledgments

We thank the Icahn School of Medicine at Mount Sinai colleagues Jeremiah J. Faith and Ilaria Mogno for helping with the characterization of the adult FMT fecal samples. This work was supported by grant no. R35 GM139655 (G.F.) from the National Institutes of Health and seed fund (G.F.) from the Icahn School of Medicine at Mount Sinai. L.E.D. lab is supported by grant no. RGPIN-2022-03915. This work was also supported in part through the computational resources and staff expertise provided by the Department of Scientific Computing at the Icahn School of Medicine at Mount Sinai.

## Author contributions

M.N. and G.F. designed the methods. M.N. performed most of the computational analysis. Y.F., M.K. and A.T. assisted with computational analysis. M.N., Y.Liu, Y. Li, W.Q., L.E.D., X-S.Z, M.K. and E.A.M. performed experiments. M.N., Y.F., Y.Liu, Y. Li, W.Q., L.E.D., X-S.Z, M.K., E.A.M., W.J., M.J.B., R.H.V. and G.F. contributed to data interpretation. M.N. and G.F. wrote the manuscript with input and comments from all coauthors. G.F. conceived and supervised the project.

## Materials and methods

### Strains, media, and growth conditions

*Escherichia coli* str. K-12 substr. MG1655 was used for heterologous expression of the gene of interest. *E. coli* strains were cultured in LB broth (LB broth Miller, Becton) or LB agar (LB agar Miller, Becton) at 37°C. When grown in liquid medium, the cultures were aerated with shaking at 220 rpm, unless otherwise specified.

*Akkermansia muciniphila* strains employed in this work include Akk_ECAM and Akk_probiotics. The Akk_ECAM strain was isolated in this study (details provided below), while the Akk_probiotics strain was derived from a commercial probiotic product (Glucose Control, Pendulum). All *Akkermansia* strains were routinely grown statically under an anaerobic condition (5% hydrogen, 5% carbon dioxide, and 90% nitrogen) using a type C anaerobic chamber with Indicator (#7250000, Coy laboratory) in BHI media (Brain Heart Infusion Agar, Gibco) supplemented with 0.25% v/v porcine gastric mucin (M1778, Sigma-Aldrich). The minimum inhibitory concentration (MIC) for amoxicillin was determined using an ETEST assay. A gradient of diluted concentrations (0.5–16 mg/L) was tested. For the selection of amoxicillin (AMX)-tolerant Akkermansia, amoxicillin sodium salt was used at a concentration of 2 mg/L.

### Stool specimen collection

ECAM samples were obtained from a previous study with an Institutional Review Board– approved study which was conducted on infants of healthy, pregnant mothers in New York City from 2011 to 2014.^1^ FMT samples were obtained from a previous study with Institutional Review Board approval.^2^

### Isolation of Akkermansia strains from infant stool

The *A. muciniphila* strain Akk_ECAM was isolated using a modified version of a previously described protocol^3^ based on growth in mucin medium (3 mM KH_2_PO_4_, 3 mM Na_2_HPO_4_, 5.6 mM NH_4_Cl, 1 mM MgCl_2_, 1 mM Na_2_S· 9H_2_O, 47 mM NaHCO_3_, 1 mM CaCl_2_, 40 mM HCl, trace elements and vitamins, and 0.25% porcine gastric mucin). To enrich for *A. muciniphila*, a loopful of stool sample (∼20 mg) was added into 2 mL of mucin medium and incubated at 37°C for 48 h under anaerobic conditions. The culture was passaged in mucin media three times by inoculating 100 μL into 2 mL of mucin medium and incubating until turbid. After three sequential passages, a suspension sample was streaked on BHI agar supplemented with 0.25% mucin and incubated for one week. Single colonies were re-streaked on BHI to purify, and the resulting purified colonies were then grown in 2 mL of mucin medium. To identify the isolated bacteria, DNA was isolated from 1 mL of culture (DNeasy Blood and Tissue Kit, Qiagen), and a portion of the 16S rRNA gene was amplified by PCR and sequenced (see primer sequences listed in **Table S6**).

### DNA extraction

#### Extraction of DNA from fecal samples

Fecal DNA was extracted with a QIAamp PowerFecal Pro DNA kit (#51804, Qiagen) according to the manufacturer’s instructions. Briefly, 50-100 mg of fecal sample was first homogenized by beads-beating using a Retsch mixer mill (MM400, Verder Scientific Inc., Newtown, PA) for 10 min at 25 Hz speed (5 min each side) and then subjected to column-based extraction steps as described in the kit protocol.

#### Extraction of DNA from bacterial cells

Genomic DNA was extracted from bacterial cells using either a Wizard Genomic DNA Purification Kit (A1120, Promega) or a Qiagen DNeasy Blood and Tissue kit (#69504, Qiagen) following the manufacturer’s instruction. The DNA was eluted in 100 µL elution buffer and stored at -20 °C.

### RNA extraction

The bacterial cell pellets were collected from 0.5 mL of cell cultures grown in BHI media supplemented with 0.25% mucin after 48h of incubation and saved in -80 °C. Total RNA was extracted with an RNeasy Mini kit (#74104, Qiagen). Briefly, cells were first lysed in RLT buffer (from the kit) with the addition of 10 μL/mL β-mercaptoethanol. The lysate was further processed by adding the content of a PowerBead tube (glass, 0.1 mm; Qiagen) and homogenizing the mixture for 5 min at a speed of 30 Hz using a Retsch mixer mill. Subsequent steps followed the manufacturer’s instructions. Contaminating genomic DNA was depleted using on-column DNase treatments with DNase (#79254, Qiagen). RNA quality was assessed with a Qubit 4 Fluorometer (Invitrogen) using the Qubit 1× dsDNA HS Assay Kit to detect DNA contamination and the Qubit RNA HS Assay Kit to measure total RNA. Ribosomal RNA was depleted using NEBNext rRNA Depletion Kit (Bacteria) (#E7850, NEB). The quality of total RNA was checked using an Agilent 2100 Bioanalyzer (#5067-4626, Agilent Technologies USA).

### Real-time quantitative RT-PCR

One microgram of total RNA was converted to cDNA using SuperScript™ VILO™ MasterMix kit (#11755050, Invitrogen). Gene-specific primer pairs are provided in **Table S6**. SYBR-based RT-qPCR was performed in 384-well plates (#4306737; Applied Biosystems) and analyzed by a qPCR system (QuantStudio 6 Flex, Thermo Fisher Scientific). The thermal cycling protocol was as follows: 10 min at 50°C, 1 min at 90°C, followed by 40 cycles of 30 s at 60°C and 1 min at 90°C, with a final melt curve stage. SYBR signal was normalized to the ROX reference dye. Cycle of quantification (Cq) values were automatically determined using QuantStudio Design & Analysis Software (v1.3). Based on the melting curves, the primers were specifically bound to the target genome. All the samples were run in replicates technical replicates.

### Library preparation and sequencing

#### Oxford Nanopore whole genome sequencing

gDNA samples were sheared to an average length of 10 kb using g-Tubes (#520079, Covaris) and purified with 0.4 × AMPure XP beads (A63882, Beckman Coulter Inc.) to remove short fragments. Native libraries were prepared using Oxford Nanopore Technologies (ONT) ligation sequencing kit and native barcoding kit (SQK-LSK109, and SQK-NBD112-24, respectively).

PCR barcoding was performed using the Native Barcoding Expansion 1-12 kit (EXP-NBD104) and/or 13-24 kit (EXP-NBD114) for sample multiplexing. Whole genome amplification (WGA) were conducted with the REPLI-g Mini kit (#150023, Qiagen) using 10 ng of input DNA and 16 h of incubation. WGA samples were treated with T7 endonuclease I (M0302S, NEB) to maximize sequencing yield, as per ONT documentation. Native or amplified gDNA samples were sequenced on R9.4.1 and R10.4.1 MinION flow cells using MinION MK1B or MK1C instruments. Basecalling was performed with Guppy v.5.0.7 (ONT) in GPU mode using the dna_r9.4.1_450bps_sup.cfg model for R9.4 data or Dorado v0.4.1 (ONT) with dna_r10.4.1_e8.2_400bps_sup.cfg model for R10.4 data.

### ONT targeted sequencing

The RMS region was amplified using specific forward and reverse primers (**Table S6**) to generate a 6.6-kb amplicon, which included sequences enable sample barcoding. PCR was performed in a 50 μL reactions containing 250 ng of gDNA and 0.7 μM of primers. Specific barcodes were assigned to the PCR products using the PCR Barcoding Expansion 1–96 kit (EXP-PBC096, ONT). A second-round PCR was performed with 0.5 nM purified PCR product, 1 × LongAmp PCR buffer, 2 mM of MgCl_2_, 0.3 mM of dNTPs and 5 units of LongAmp Taq DNA polymerase (NEB) in a 50 μL reaction. The thermal cycling protocol was 95°C for 3 min, followed by 15 cycles of 95°C for 15 s, 62°C for 15 s, 65°C for 7 min, with a final extension at 65°C for 7 min. Amplicons were sequenced on R9.4.1 / R10.4.1 Flongle flow cells.

### PacBio WGS sequencing

The SMRTbell library was prepared according to the manufacturer’s instructions. DNA was sheared to ∼5,000 bp using a Covaris LE220-plus Focused-ultrasonicator (#500569, Covaris). The samples underwent damage and end repair, followed by ligation of single-stranded barcoded adaptors. Sequencing primer was annealed to the SMRTbells and incubated with templates for 1 hour at 20 °C using a primer-to-template ratio of 20:1. The Sequel 2.1 polymerase was bound to the annealed complex at a polymerase to SMRTbell ratio of 30:1 and incubated for 4 h at 30 °C. The binding complex was cleaned, diluted to the final loading concentration (125-175 pM), and held at 4 °C until diffusion-based loading on the Sequel II instrument (PacBio). The sequencing runs used 8M sequencing chips, with 30 h movies recorded following the manufacturer’s guidelines. Most metagenome samples except for some FMT samples, were sequenced on the latest Revio instrument (PacBio). Circular consensus sequencing (CCS) reads were generated using CCS v.3.4.1 (https://github.com/PacificBiosciences/ccs) with default settings.

### Illumina RNA sequencing

RNA-seq library construction for Illumina sequencing was performed using the NEBNext Ultra II RNA Library Prep Kit for Illumina (#E7770, NEB). Library quality and size were assessed using an Agilent 4200 TapeStation system (Agilent Technologies, Inc.) and quantified with a Qubit 4 Fluorimeter using Qubit DNA HS Assay Kit (Invitrogen). All samples were equitably pooled, size-selected, and sequenced on an Illumina HiSeq 2500 platform, generating paired-end reads of 150 base pairs (bp) each.

### Phylogenetic analysis of HMP reference genomes

The bacterial genomics sequences and proteome files were downloaded from the NIH Human Microbiome Project (HMP) legacy data browser website (https://www.hmpdacc.org/hmp/HMRGD/). A phylogenetic tree was constructed using PhyloPhlAn v3.0.67^4^ with proteome files as input and high--diversity configuration. The tree was visualized and edited using the interactive tree of life (iTOL) software (https://itol.embl.de).

### *De novo* assembly of metagenomes

#### ONT read metagenomic assembly and taxonomic annotation

Basecalled reads were processed using metaFlye tool v2.9^5^ with the parameters ---nano-raw -- meta --keep-haplotypes --iterations 2. Meta-assemblies were divided into species-level bins using the MaxBin 2.0 software.^6^ Strainberry v1.1^7^ was used to phase haplotypes and separate strain level contigs. Contigs were subsequently binned using the "binning" module of metaWRAP v1.3.2.^8^ The quality of the reconstructed MAGs was evaluated using the ’bin_refinement’ module of metaWRAP v1.3.2, and taxonomic assignment was performed using GTDB-tk v3.8.^9^

#### SMRT read metagenomic assembly and taxonomic annotation

HiFi sequencing reads were assembled using hifiasm_meta 0.13-r308 (https://github.com/xfengnefx/hifiasm-meta) with default parameters, focusing on primary contigs. To maximize MAGs recovery, the HiFi-MAG-Pipeline (https://github.com/PacificBiosciences/pb-metagenomics-tools/tree/master/HiFi-MAG-Pipeline) was deployed, increasing both the total number of MAGs and single-contig circular MAGs.

#### Hybrid assembly

Nanopore-based long reads and Illumina paired-end short reads were used for a hybrid assembly with MUFFIN,^10^ producing high-quality and complete bins.

### Gene annotation and RMS prediction

Gene annotation for bacterial genomes and MAGs was performed using Prokka v1.14.6.^11^ RMS identification followed previously established methods.^12^ Curated reference protein sequences for types I, II, IIC, and III RMS, and type IV REases were downloaded from the REBASE^13^ “gold standards” dataset (accessed November 2016). A dedicated BLAST database was created for the specific gene (HsdS) of Type I RMS using 39,729 HsdS proteins from REBASE. The TRD domains in hsdS genes were identified using HMMER v3.3 (http://hmmer.org) with a profile built from 31 TRD domains.

### Detection of SNPs and small Indels in antibiotic-treated *A. muciniphila* strains

Genome variations were identified from the aligned Nanopore reads to the reference genome using Clair3^14^ with default ONT settings, except for the options --no_phasing_for_fa, -- include_all_ctgs, and --haploid_precise. Variants called by Clair3 were filtered with bcftools^15^ based on the criteria "QUAL > 15 and DP > 10". Subsequently, SNPs located within structural variation regions were excluded. The potential effects of the candidate SNPs were then annotated using snpEFF.^16^

### Differential-expression analysis of RNA-seq data

Read quality was assessed using FastQC v.0.11.9 (http://www.bioinformatics.babraham.ac.uk/projects/fastqc). Adapter sequences and low-quality reads were removed using Trimmomatic v.0.36^17^ with the following parameters: PE, -phred33, ILLUMINACLIP:<adapters.fa>:2:30:10:8: keepBothReads, SLIDINGWINDOW:4:15, LEADING:20, TRAILING:20, and MINLEN:50. rRNA sequences were then filtered out using SortMeRNA v.2.1b ^18^ with the SILVA 16S/23S^19^ and the Rfam 5S^20^ rRNA database. Non-rRNA reads were aligned to the reference genome using BWA-MEM v.0.7.17-r1198.^21^ Read assignment was performed with FeatureCounts v.2.0.1,^22^ excluding multi-mapping and multi-overlapping reads. Normalization and differential expression testing were conducted using DESeq2 v.1.18.1,^23^ with differentially expressed genes defined as those with an FDR-corrected *p* < 0.05 and |log2[FC]| > 1.

### Gene set enrichment analysis

Gene function was annotated by eggmapper v2.1.6 against the defaults eggNOG database.^24^ GO and KEGG enrichment analyses were performed using the R package clusterProfiler v.3.18.1. ^25^ KEGG pathways and modules were filtered for UniRef90 v.2019_06 entries, downloaded from UniProt website.^26^ Enrichments at different taxonomic levels were calculated and visualized as heatmap using ComplexHeatmap package v.2.6.2.^27^

### Genome-wide epigenetic phase variation (ePV) detection in long-read data

#### Structural variation (SV) detection

Basecalled ONT reads were aligned to the de novo assembled MAGs with NGMLR v 0.2.6.^28^ HiFi PacBio reads were aligned to the de novo assemblies with pbalign (SMRTlink v10, https://downloads.pacbcloud.com/public/software/installers/smrtlink_10.1.0.119588.zip). The resulting alignment files were processed with sniffles v2.0.7^29^ for structural variant calling. Detected SVs were mapped to their corresponding genome locations, and SVs overlapping predicted RMS regions / orphan MTases (within a 2,000 bp window) were defined as potential ePVs. The ePV list was refined through manually curation.

#### Quantification of ePV with raw reads

To overcome the limitations of SV detection tools in quantifying read counts assigned to genomic variants in complex rearrangement regions, we developed an in-house pipeline: (1) predicted all possible genomic variants base on the reference genome and the read alignment; (2) created an *in silico* reference for each variants and merged them; (3) realigned reads to the new reference genomes; (4) applied strict filtered thresholds, including “MAPQ 60 & mg > 99” and “-max softsclip 300”, (5) counted the reads uniquely mapped to each genotype.

### Site-specific ePV detection for PacBio data

#### Preprocess of PacBio raw data

PacBio sequencing data contained both DNA sequence and kinetic data, which were used to detect single-molecule methylation signal. Raw SMRT Sequel II data was processed into CCS reads and demultiplexed using SMRTlink v10.

#### Genome-wide aggregated methylation analysis

Subreads and CCS reads were aligned to MAGs using pbalign (SMRTlink v10) with default parameters, and alignments were separated by MAG. The IPD ratio was calculated with ipdSummary (SMRTlink v10) for each MAGs. De novo methylation motifs were identified using motifMaker (SMRTlink v10) with default parameters. All of the above analyses aggregate the CCS data based on the MAG genomes, as opposed to single-molecule analysis, which is based on each individual CCS reads.

#### Single molecule level methylation analysis

Subreads were split based on CCS read ID and mapped to their corresponding CCS reads using pbalign with default parameters. Alignments were filtered with strict thresholds, including “MAPQ 60 & mg > 99” and “-max softsclip 300”. Single-molecule IPD ratio was calculated with ipdSummary. For Revio data, raw data lacked subread information and require an in-house scripts for single-molecule analysis (released the detailed scripts on Zenodo).

#### Identification of single site ePV

IPD ratio files from single-molecule methylation signals were processed in R v4.0.3. Low methylation sites (< 50%) were identified on both strands, and only sites with low methylation on both strands were considered partially methylated sites. The methylation variation sites were visualized across MAGs, with loci and potentially affected genes highlighted in a gene diagram chart.

#### Genome-wide ePV detection in Illumina short-read data

For short-read metagenomic data, ePVs exclusively referred to invertons, as defined in a previous study.^30^ Invertons across the metagenomic samples were identified using the PhaseFinder^30^ pipeline, with HMP reference genomes (https://hmpdacc.org/hmp/HMRGD/) as reference. Invertons overlapping predicted RMS regions / orphan MTase (within a 2,000 bp window) were classified as potential ePVs.

### Protein extraction, purification and identification

#### Protein preparation

*E. coli* MG1655 cells were harvested by centrifugation at 5,000 × g for 10 minutes at 4°C. The supernatant was collected in 50 mL aliquots. The cell pellet was resuspended in lysis buffer containing 50 mM Tris-HCl (pH 7.5), 150 mM NaCl, 2 mM tris (2-carboxyethyl)phosphine (TCEP, #68957, Sigma-Aldrich) and a protein inhibitor cocktail (P8340, Sigma-Aldrich). The suspension was lysed by sonication on ice (15-second pulses, 10 times, with 2-min intervals). The lysate was clarified by centrifugation at 12,000 × g for 30 min at 4°C to remove cell debris. The supernatant from cell culture was concentrated to 1 mL using an Amicon Ultra Centrifugal Filter unit (3 kDa, Millipore) and subjected to sonication as described above. The resulting crude protein solution was purified using a Capturem His-Tagged Purification kit (#635719, TaKaRa Bio Inc.) following the manufacturer’s instructions.

#### SDS-PAGE

Protein samples were mixed with 4× Sodium dodecyl sulfate (SDS) loading buffer (Bio-Rad Laboratories, Inc) and heated at 98°C for 5 min. SDS-polyacrylamide gel electrophoresis (SDS-PAGE) was performed using Mini-PROTEIN TGX Gels (4–20%, Bio-Rad Laboratories, Inc). Gels were stained with Coomassie Brilliant Blue R-250, and protein bands were visualized using a gel documentation system (Bio-Rad Laboratories, Inc).

#### Intact Protein Mass Spectrometry

For intact protein mass analysis, purified protein samples were buffer exchanged into 1 M ammonium acetate solution using a 3 kDa Amicon Ultra Centrifugal Filter unit (Millipore). Qualitative analysis of recombinant protein was performed with a 1260 HPLC system coupled to a 6545 XT Q-TOF MS instrument operated in positive ESI mode (Agilent Technologies, Inc.).

The acquired spectra were extracted and deconvoluted with Bioconfirm A 2.0 software (Agilent Technologies) with the default parameters, except that the mass range was set to be 10 kDa to 100 kDa.

### In vitro passage assay of *A. muciniphila*

#### AMX selection assay

An *A. muciniphila* isolate with the g1 RMS genotype was selected as the starting material and cultured on BHI agar plates supplemented with 0.25% mucin for three days. 46 single colonies were picked and subcultured in 3 mL BHI broth supplemented with 0.25% mucin for two additional passages. The above subcultures were diluted to an initial OD_600_ of 0.01 and a 50 μL aliquot was inoculated in 150 μL BHI liquid medium supplemented with 0.25% mucin, and with or without 2 mg / L AMX. The subculture process was repeated for nine passages. Cells from each passage were collected and stored. gDNA extraction and targeted sequencing were performed as described above.

#### RMS Reversible assay

To assess the conditions under which the RMS genotype could revert from non-g1 to g1, *A. muciniphila* colonies with the non-g1 RMS genotype from the AMX selection assay were stressed with various microbiological mediums, antibiotics, and atmospheric conditions (see detailed design and conditions in **Table S7**). RMS genotypes were determined across the assay using targeted sequencing. Additional replicates for ciprofloxacin treatment were performed following the same procedure but with an increased number of replicates. The minimum inhibitory concentration (MIC) for each antibiotic was determined using a preliminary CFU count assay. Antibiotic concentrations were selected based on their respective MIC values.

### Expression of the *mucC* gene in *E. coli*

To overexpress the *mucC* gene, we synthesized the gene with codons optimized for expression in *E. coli*. The gene was inserted into a medium-copy-number plasmid containing a constitutive expression promoter. The recombination plasmid was then transformed into *E. coli*. Overnight cell cultures were diluted 1:100 into fresh LB medium containing different concentrations of AMX. The growth curve of each group of cells (OD_600_) was monitored using a microplate spectrophotometer (SpectraMax M5, USA).

#### AMX sensitivity test

AMX test strips (AES, Liofilchem) were placed on the surface of a BHI agar plate supplemented with 0.25% mucin and inoculated with a 500 μL bacterial suspension. The plates were incubated at 37°C for 48 h under anaerobic conditions. The MIC was determined by the intersection of the lower part of the ellipse-shaped growth inhibition area with the test strip.

### Akkermansia-fed Mouse model

A probiotic supplement including *Clostridium beijerinckii* WB-STR-0005, *Clostribium butyricum* WB-STR-0006, *A. muciniphila* WB-STR-0001 (introduced earlier as Akk_probiotics), *Anaerobutyricum hallii* WB-STR-0008, and *Bifidobacterium infantis* 100 (Pendulum, USA) was administered to healthy male germ-free (GF) C57BL/6J mice (JAX stock #000664) (n=4). 12-week-old GF mice were fed 10 mg probiotic supplement via Mini Capsule Kit (Braintree Scientific XKITM) for the first two days. Fecal samples were collected at day 7.

### Quantification and statistical analysis

Statistical analyses were performed using R (v.4.0.3). Statistical significance was determined using unpaired two-tailed Mann–Whitney–Wilcoxon test, unpaired two-tailed Welch’s t-test, and unpaired Student’s t-test, as indicated in figure legends. Data are shown as mean ± s.d. *P* < 0.05 was considered statistically significant. Information regarding the number of samples and statistical tests are provided in the figure legends.

