## Supplementary Figures for "Epigenetic phase variation in the gut microbiome enhances bacterial adaptation"

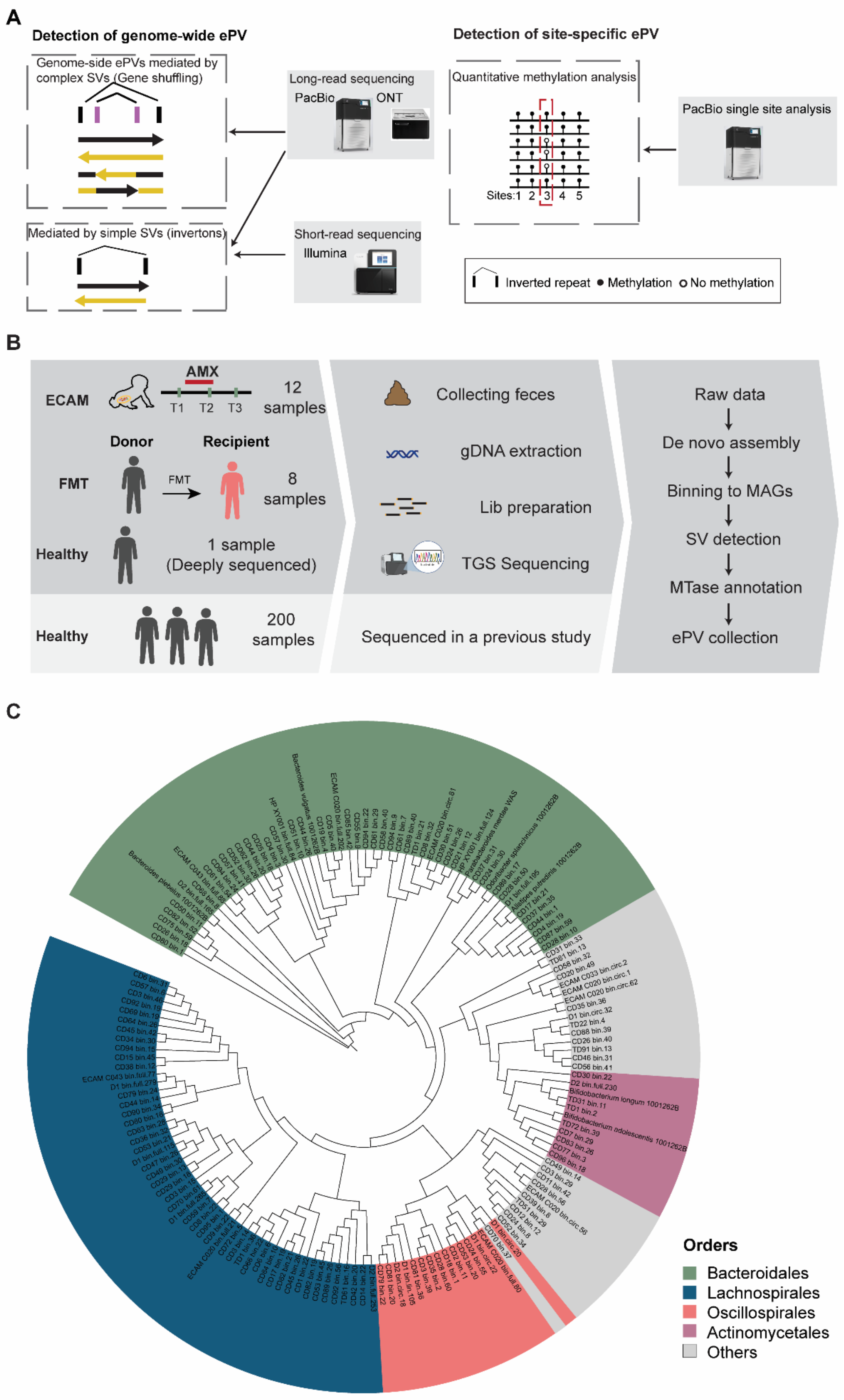
A
Detection of genome-wide ePV
Genome-side ePVs mediated by complex SVs (Gene shuffling)
Mediated by simple SVs (invertons)
Long-read sequencing
PacBio
ONT
Short-read sequencing
Illumina
Detection of site-specific ePV
Quantitative methylation analysis
Sites:1 2 3 4 5
PacBio single site analysis
Inverted repeat
Methylation
No methylation
B
ECAM
AMX
T1 T2 T3
12 samples
Donor
Recipient
FMT
FMT
8 samples
Healthy
1 sample (Deeply sequenced)
Healthy
200 samples
Collecting feces
gDNA extraction
Lib preparation
TGS Sequencing
Sequenced in a previous study
Raw data
De novo assembly
Binning to MAGs
SV detection
MTase annotation
ePV collection
C
Orders
Bacteroidales
Lachnospirales
Oscillospirales
Actinomycetales
Others

**Figure S1. Comprehensive detection of genome-wide ePVs across taxa using long-read metagenomics**

(**A**) Comprehensive analysis of genome-wide (Left panel) and site-specific (Right panel) ePVs by both long- and short-read sequencing. (**B**) Workflow of detecting and profiling genomic ePV from the gut microbiome with long-read sequencing technology. In this study, we sequenced 20 samples from the ECAM (infant exposed to antibiotics)[1] and FMT (adult *Clostridioides difficile* infection patient) cohorts. Additionally, we performed deep sequencing on a healthy adult sample. We also use data from 200 healthy adult samples previously sequenced in another study.[2] TGS, Third generation sequencing. (**C**) Phylogenetic analysis of strains with ePVs. Maximum likelihood phylogenetic tree was constructed for 158 MAGs based on the alignment of 399 marker genes using PhyloPhlAn.[3] The phylogenetic tree is color-coded by order level.

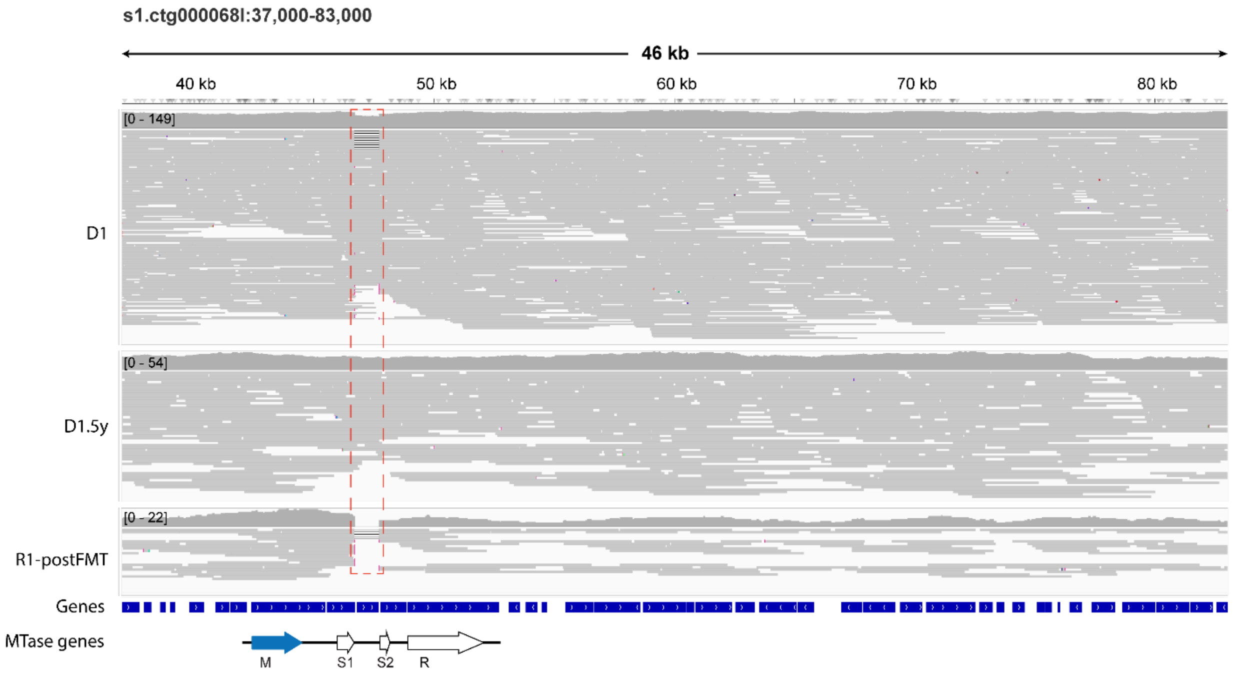


**Figure S2. Reliable detection of intra-strain variation in the human gut microbiome**

Alignment tracks for the *G. qucibialis* strain in the FMT donor samples (donor: D1; donor 5 years: D1.5y ) and the recipient sample one year after FMT (R1-postFMT) are displayed in the corresponding IGV screenshot. Apart from the genome-wide ePV (highlighted with a red dashed box), no significant SNPs were detected across the three samples, confirming that the same strain was analyzed. Genes, including MTase-related genes, are annotated in dedicated tracks. M: MTase; S1, specificity unit 1; S2, specificity unit 2, R: REase.

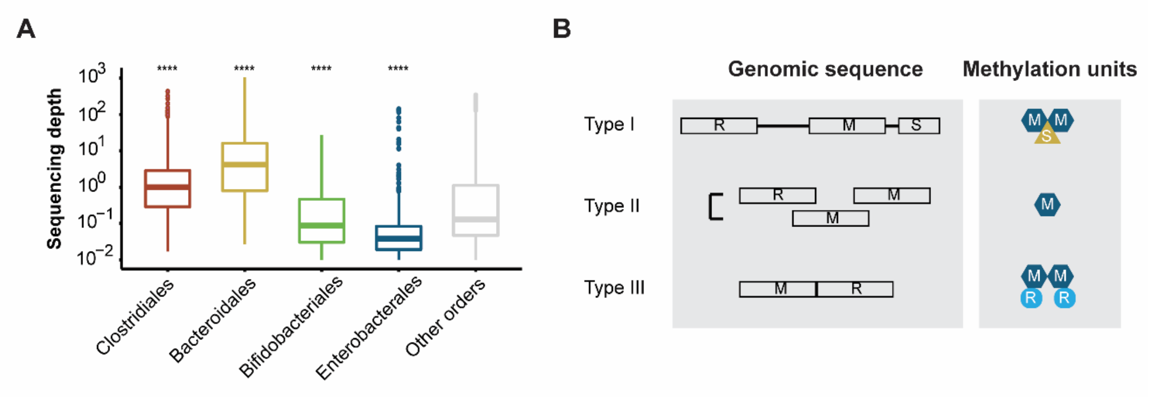


**Figure S3. Comprehensive detection of genome-wide ePVs using published NGS metagenomics**

(**A**) Sequencing depth per bacterial strain in HMP metagenomic samples. Alignment files for the HMP samples were obtained from the HMP portal, and sequencing depth per genome was calculated and summarized at the order level. Significance levels are indicated as * p ≤ 0.05, ** p ≤ 0.01, *** p ≤ 0.001, **** p ≤ 0.0001. (**B**) Types of restriction-modification system in bacteria. R: Restriction enzyme, M: MTase, S: Specificity unit.

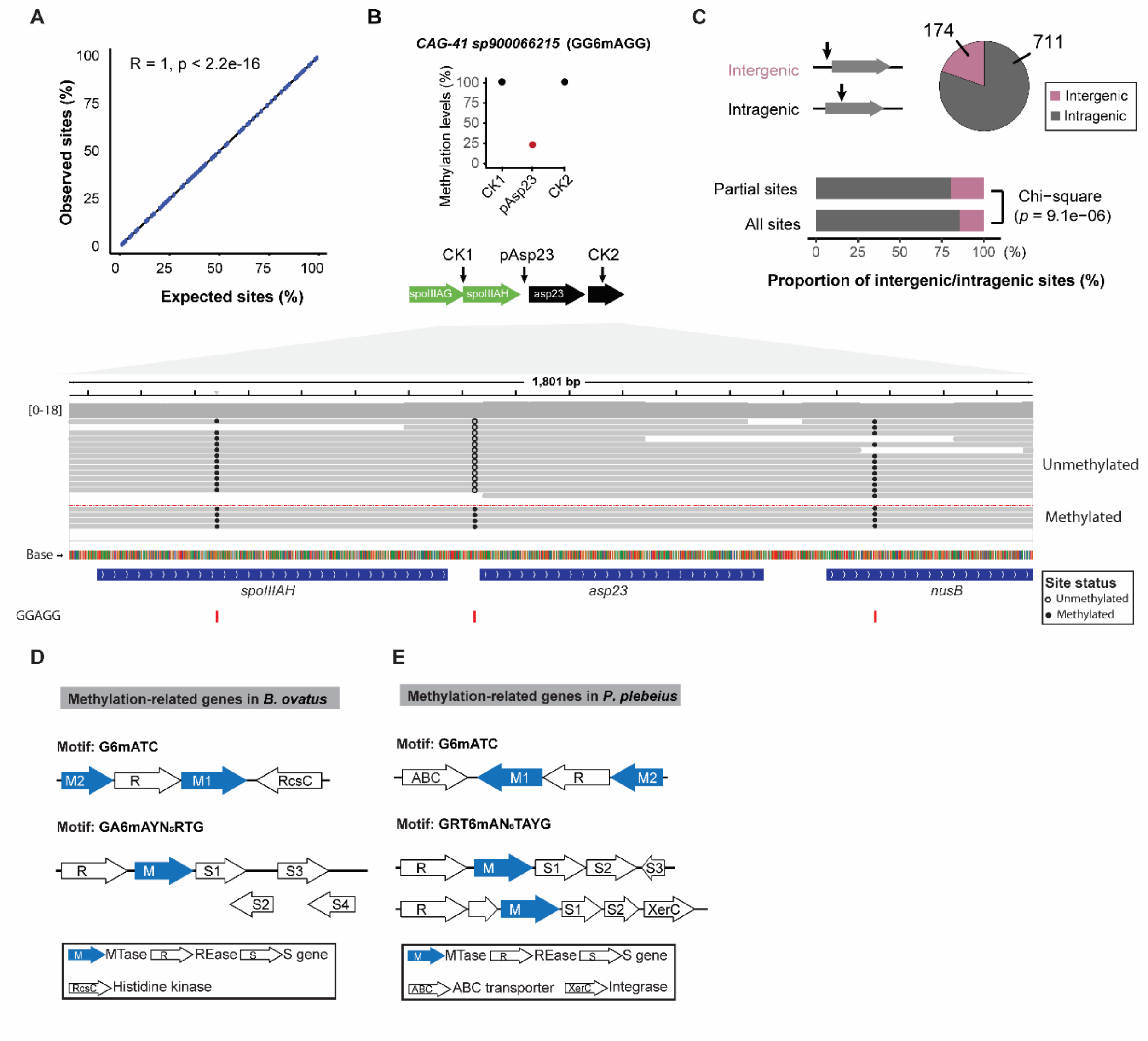


### Figure S4. Comprehensive detection of site-specific ePVs in the human gut microbiome

(**A**) Simulations of GATC site mixtures were conducted using fully methylated (*E. coli* K12 wild type) and methylation-free (*E. coli* K12 ER3413) SMRT data, encompassing varying levels of methylation. Single-site methylation analysis was performed using an in-house developed pipeline. The methylation level at each G6mATC site was quantified, and the correlation between expected and observed values was subsequently evaluated. (**B**) A partially methylated GG6mAGG site was observed in the CAG-41 *sp900066215* strain from Infant 2. The summarized methylation level is presented in the point plot. The methylation status of each molecule at this specific site, located in the promoter region of asp23 (pAsp23), is shown in the IGV screenshot below which illustrates the per-read methylation status across the three investigated sites. Two neighboring GG6mAGG sites were used as control sites. All three GGAGG sites are marked with arrows in the gene diagram. (**C**) Genetic types of site-specific ePV sites. Chi-square test was performed to compare the proportions of intergenic sites in partially methylated sites versus all methylation sites. (**D**) Methylation-related genes and methylation motifs observed in a *Bacteroides ovatus* strain from Infant 3 in the ECAM cohort. (**E**) Methylation-related genes and methylation motifs observed in a *Phocaeicola plebeius* strain in the FMT cohort. (**D**, **E**) Only MTases with assigned methylation motifs were included. R: Restriction enzyme, M: MTase (highlighted in blue); S1, specificity unit 1; S2, specificity unit 2, R: REase.

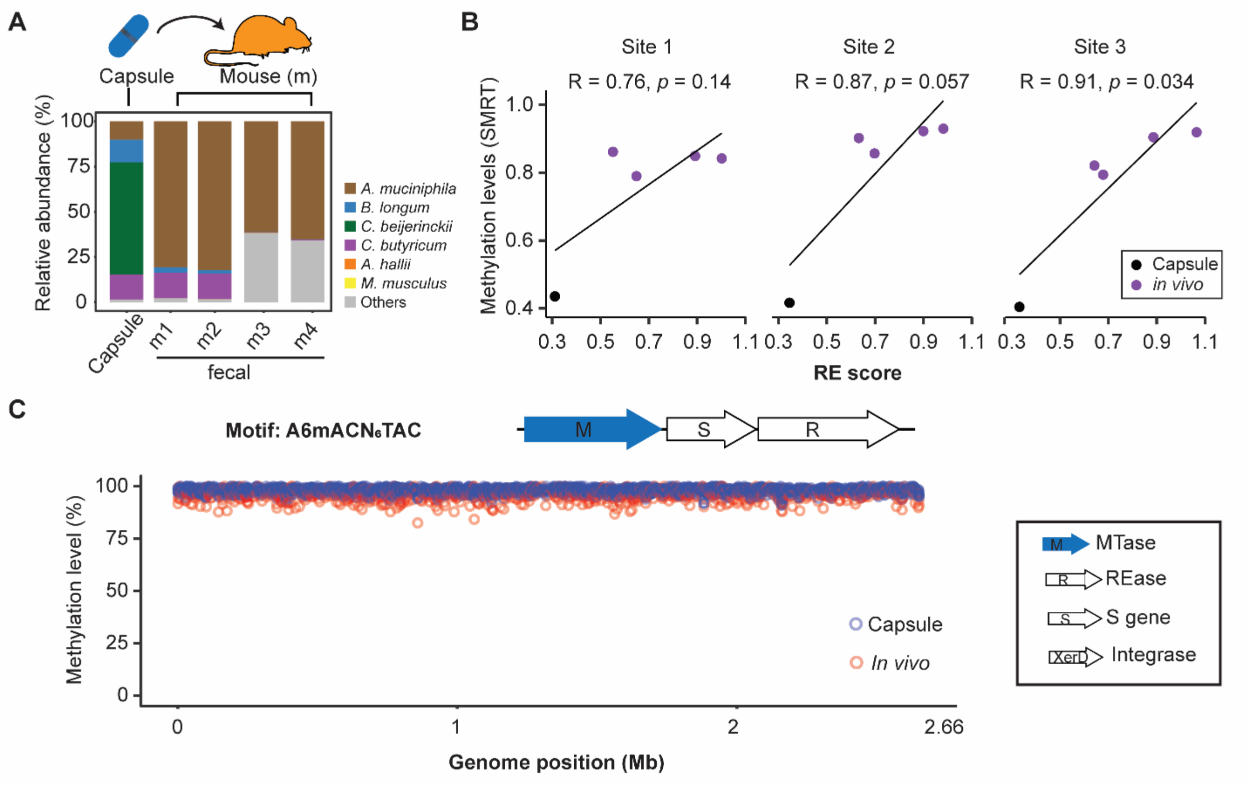

**Figure S5. Independent validation of the reliability of site-specific ePV quantification in a probiotic *Akkermansia* strain**

(**A**) Relative abundance profile of species in fecal samples from Akk-fed mice. Raw metagenomic reads were aligned to reference genomes, which included five bacterial species (see Materials and Methods) and *M. musculus*. The relative abundance was calculated as the percentage of reads obtained for a specific taxon related to the total number of reads and then normalized by genome size. (**B**) Partially methylated R6mAATTY sites involved in the colonization of the probiotic *Akkermansia* strain in mice. The three partially methylated sites identified in **Fig. 4K** were validated by restriction digestion method which specifically cut the unmethylated motif but left methylated ones intact. (**C**) Single-site methylation analysis of the A6mACN$_6$TAC motif in the probiotic *Akkermansia* strain. The methylation level of each adenine site in the A6mACN$_6$TAC motif was determined and mapped across the genome. All investigated sites were highly methylated in both capsules (highlighted in blue) and *in vivo* samples (highlighted in red). This analysis served as a negative control for estimating site-specific ePVs in the *Akkermansia* strain. R: Restriction enzyme, M: MTase (highlighted in blue), S: Specific subunit.

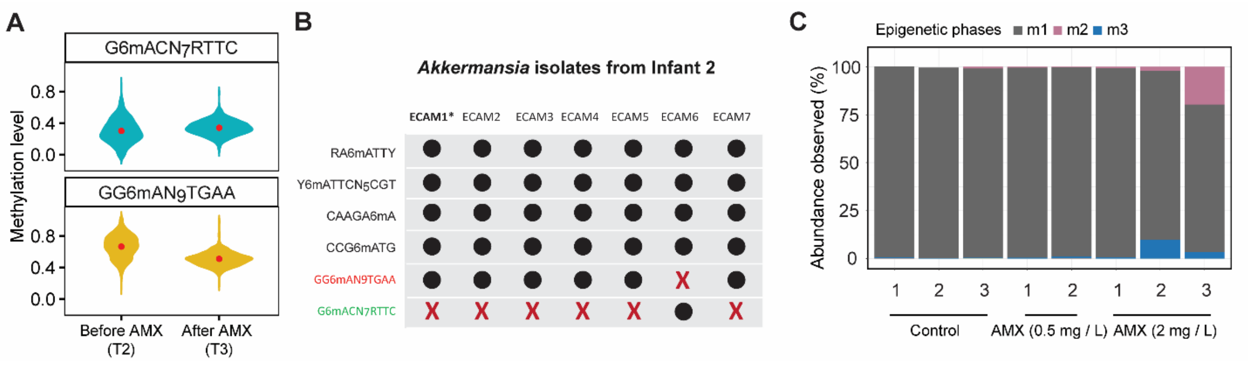


**Figure S6. Genome-wide ePV detection in both the fecal samples and *in vitro Akkermansia* isolates**

(**A**) Genome-wide methylation changes in fecal samples from infant 2 in the ECAM cohort. After AMX treatment, the subpopulation with m1 methylation (GG6mAN$_9$TGAA) decreased, while the non-m1 methylation subpopulation (G6mACN$_7$RTTC) increased. (**B**) Methylation motif profiles in *Akkermansia* isolates from fecal samples of ECAM Infant 2. All the isolates were sequenced using ONT R9 flow cells, generating matched native and WGA reads. Nanodisco[4] was utilized for methylation analysis, including base modification detection and motif calling. The motifs methylated by the phase variable MTase were highlighted in red (m1 methylation) and green (non-m1 methylation). An X-marker indicates that the motif was not detected in the corresponding isolate. ECAM1*, referred to as Akk_ECAM, is the strain used for the *in vitro* passage assay. (**C**) MTase genotype changes in Akk_ECAM under different AMX concentrations. The *A. muciniphila* strain (m1 dominant, referred to as Akk_ECAM) was exposed to AMX at concentrations of 0.5 mg/L and 2 mg/L for three consecutive passages. To assess the impact of AMX on epigenetic phase variations, targeted sequencing was performed on samples from passage 3. Only epigenetic phases with an abundance > 10% are shown.

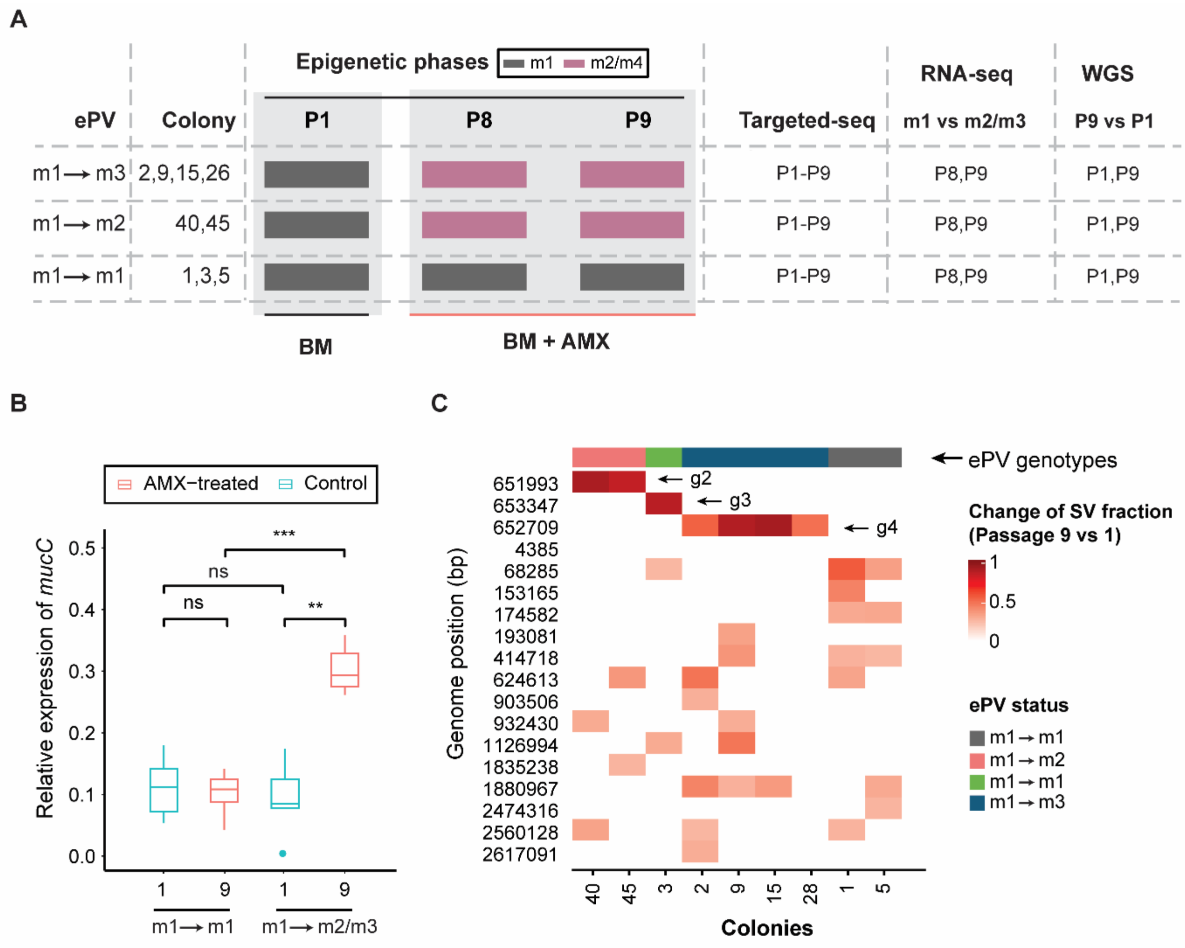


**Figure S7. RNA-seq and whole-genome sequencing reveal genotype-specific responses to AMX in the AKK_ECAM strain**

(**A**) Experimental design for RNAseq and shotgun whole genome sequencing (WGS) of samples from the *in vitro* passage assay. There are two categories of epigenetic phases, m1 and non-m1 (m2/m3), in the Akk_ECAM strain. AMX treatment is hypothesized to select for the non-m1 phase under both *in vitro* and *in vivo* conditions. We propose that the shift from m1 to non-m1 influences the expression of *A. muciniphila* genes, increasing its tolerance to AMX. To test this hypothesis, we selected a total of nine clones with either the m1 or non-m1 epigenetic phase and profiled their transcripts using RNA-seq at Passages 8 and 9 in the presence of AMX treatment. To investigate genomic changes associated with AMX exposure, gDNA of those nine colonies were also extracted at two key time points: Passage 1 (before AMX treatment) and

Passage 9 (last passages under AMX selection). Shotgun WGS was performed on an ONT R10 flow cell. (**B**) Relative expression of *mucC* in AMX-treated and control samples. Gene expression was quantified using qRT-PCR and the comparative threshold (ΔΔCt) method. *gyrA* and *dnaG* were used as reference genes for data normalization. (**C**) Structural variation (SV) detected in different colonies from *in vitro* passage assay. All the passages from each colony were sequenced using ONT R10 flow cells. Sniffle v2 was used for SV detection. SV fraction changes were calculated for each passage and visualized on a heatmap. No SV was associated with the ePV events. MTase variants were labeled with arrows.

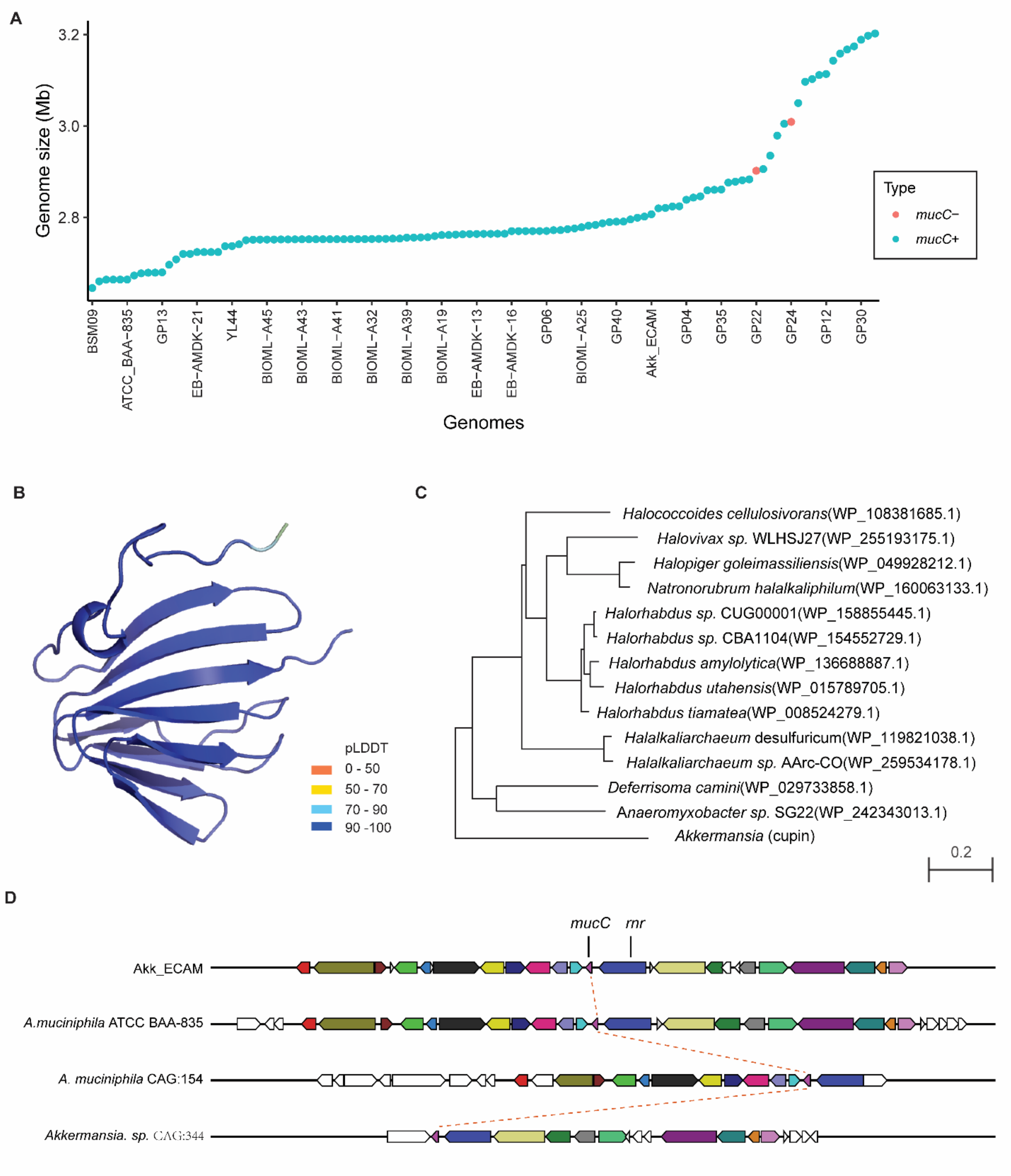


**Figure S8. Conservation and structural analysis of MucC protein in *A. muciniphila***

(**A**) Conservation of mucC in *A. muciniphila* strains. The *mucC* gene were detected in 111 out of 113 *A. muciniphila* strains (**Table S5**). Strains are highlighted in red for mucC-negative and blue for mucC-positive. (**B**) 3D structure prediction of MucC. Ribbon diagrams were colored based on per-residue pLDDT scores (range 0 to 100, with higher scores indicating better confidence). These scores were stored in the B-factor column of the .pdb; thus, when colored as pseudo-B-factors in Pymol (The PyMOL Molecular Graphics System, v 3.0 Schrödinger, LLC.), a gradient from orange to blue represents the lowest to highest pLDDT scores. (**C**) Homologs of *mucC* in other species exhibit high similarity. The amino acid sequence of MucC from the Akk_ECAM strain was used as a query to search against the NCBI nr database using Blasp program. (**D**) Genome context analysis of *mucC* across representative members of sequenced *Akkermansia* species. The cluster containing *mucC* and 12 flanking genes on each side was used as a search query for the program Multigeneblast[5], which identifies multiple gene blast hits within close genomic proximity. Homologous genes are represented with consistent coloring.

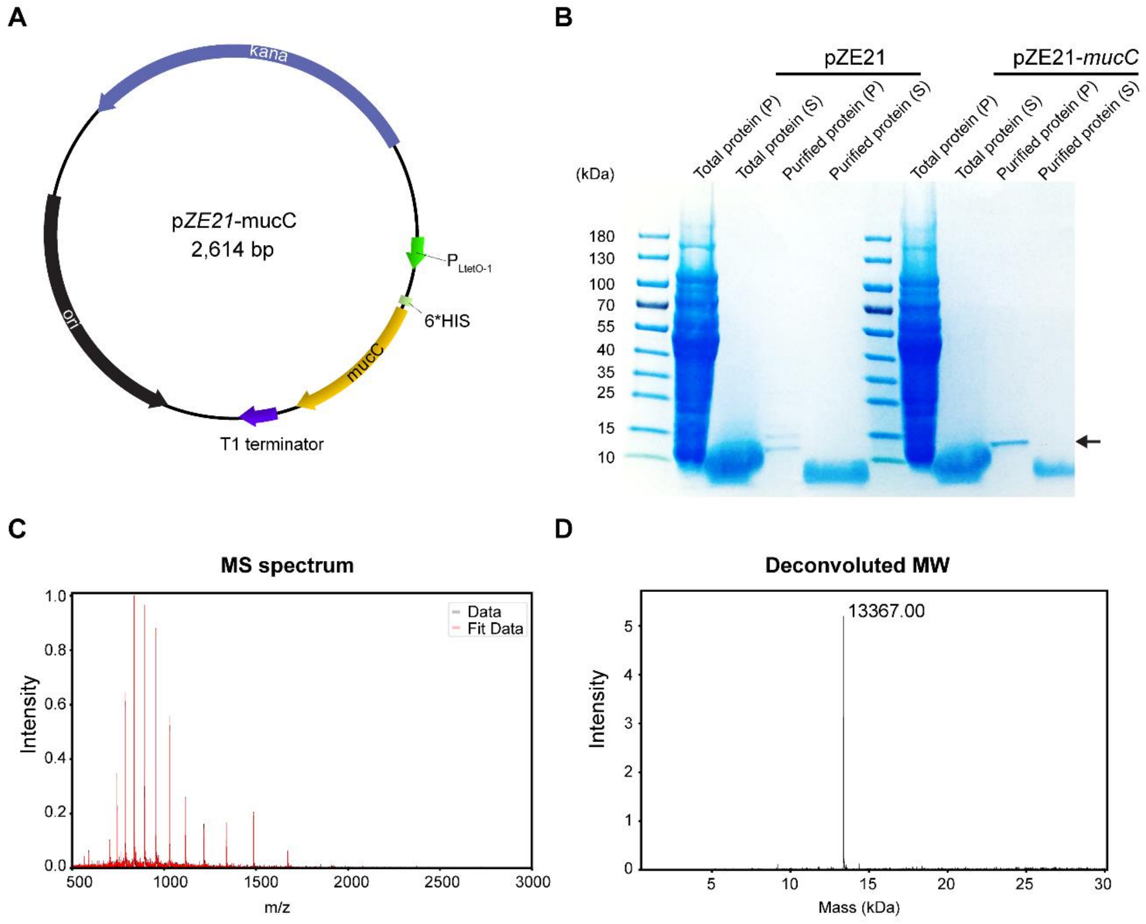


**Figure S9. Plasmid construction, expression, and validation of the *mucC* gene in *E. coli***

(**A**) Construction of a vector for overexpression of *mucC* in *E. coli* strain MG1655. The *mucC* gene was synthesized with codons optimized for expression in *E. coli*. The synthesized DNA fragment was then inserted into the pZE21 vector, resulting in plasmid pZE21-*mucC*. The expression of *mucC* was placed under the control of the LtetO-1 promoter. (**B**) The recombinant MucC protein was successfully expressed in the *E. coli* MG1655 cells. Total protein and Ni-affinity purified protein were analyzed using SDS-PAGE. P, pellet; S, supernatant. (**C**) The recombinant protein was further characterized by intact protein mass spectrometry. (**D**) Deconvoluted molecular weight analysis confirmed the identity of the recombinant MucC protein.

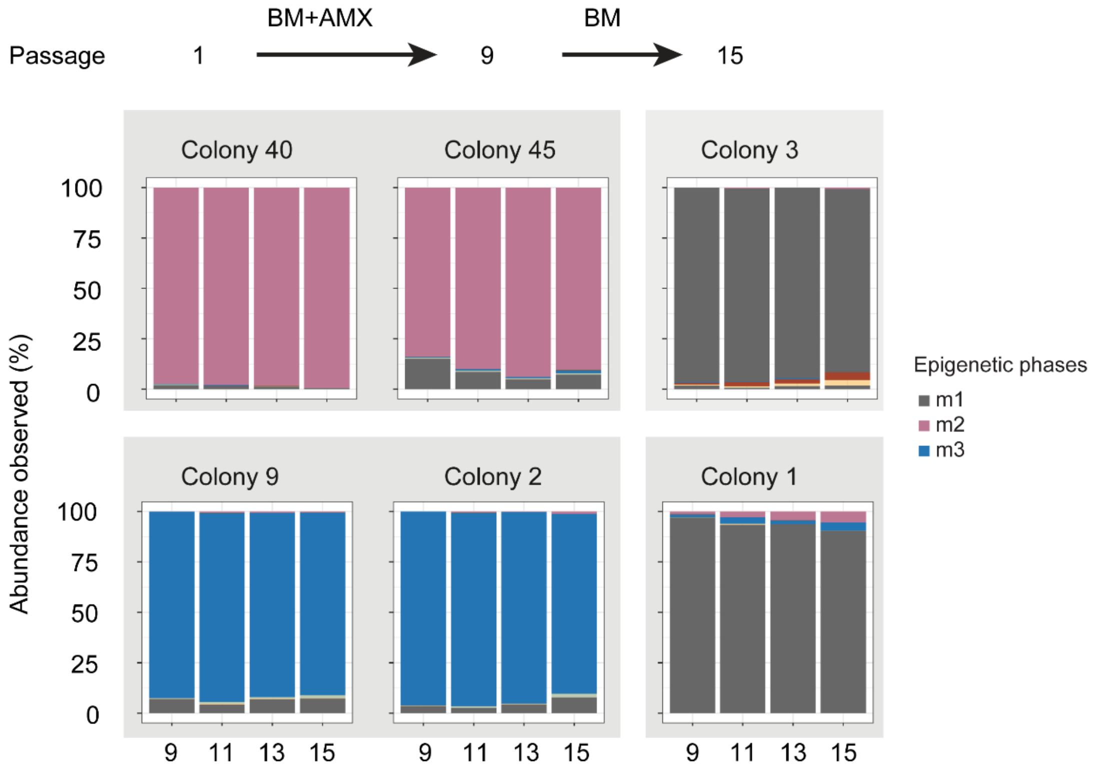


**Figure S10. Removal of AMX did not revert selection to the m1 ePV state**

To determine whether non-m1 selection imposes a fitness cost in vitro, we removed AMX at passage 10 and continued for an additional five passages. In these AMX-selected non-m1 colonies (2#, 9#, 40#, 45#), the epigenetic phase remained predominantly m2/m3 through passage 15, even in the absence of AMX treatment. Colonies 1# and 3# served as a negative control, as their epigenetic phase remained unchanged throughout passages 1 to 15, regardless of AMX treatment. Only epigenetic phases with an abundance > 10% are shown in legend

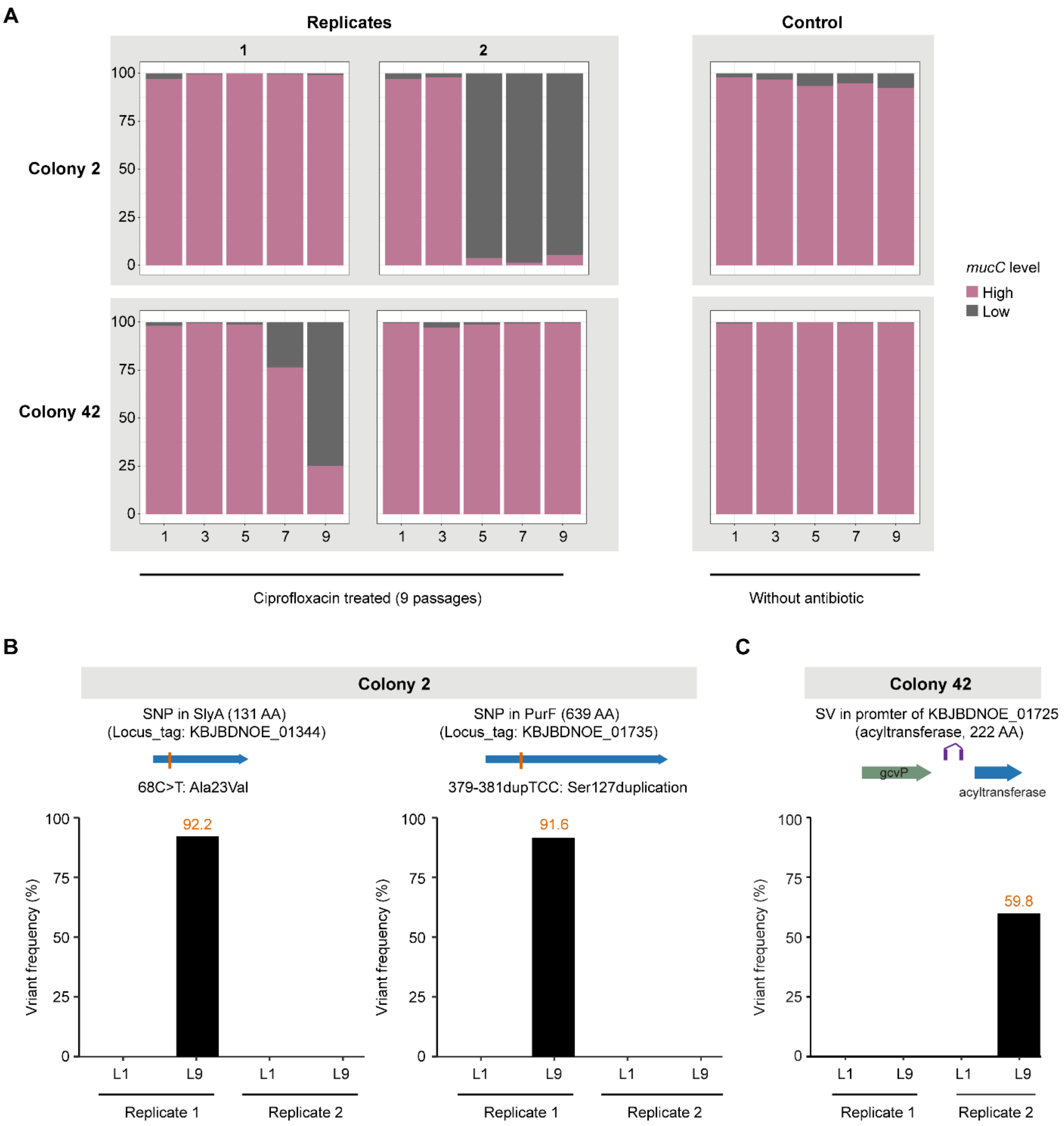


**Figure S11. Genetic variations explain the different selection outcomes by ciprofloxacin**

**(A)** Two representative clones derived from the in vitro AMX passage assay were treated with sub-lethal ciprofloxacin concentrations (50 mg/L). *mucC* expression levels, indicated by MTase genotype, were determined at every other passage (1, 3, 5, 7, 9) through targeted sequencing. Replicates for each clone and cultures without antibiotics served as negative controls. (**B**) To investigate the varying ciprofloxacin responses observed in different replicates of clone 2, shotgun sequencing was performed on passage L1 and L9 samples using the ONT R10 flow cell. For replicate 1, which did not exhibit m1 selection by ciprofloxacin, two SNPs were identified in passage L9 and their potential gene effects were predicted. The first mutation was located in the transcriptional factor SlyA (Ala23Val), potentially associated with resistance to multiple antibiotics.[6] The second mutation was found in the gene body region of PurF (Ser127duplication), as reported in a recent study.[7] (**C**) To investigate the varying ciprofloxacin responses observed in different replicates of clone 42, shotgun sequencing was performed on passage L1 and L9 samples using the ONT R10 flow cell. For replicate 2, which did not exhibit m1 selection by ciprofloxacin, a candidate structural variation (inversion) was identified in passage L9 and its potential gene effects were predicted. The inversion was located in the promoter region of an acyltransferase family protein.
