## Supplementary Data for "Epigenetic phase variation in the gut microbiome enhances bacterial adaptation"

**These IGV plots illustrate reliable detection of intra-strain variation across different species.**

Alignment tracks for the *Bacteroides xylanisolvens* (**A**), *Akkermansia muciniphila* (**B**), *Phocaeicola dorei* (**C**), *Bacteroides fragilis* (**D**), *Bifidobacterium kashiwanohense* (**E**), *Bacteroides ovatus* (**F**) strains in the ECAM samples are displayed in the corresponding IGV screenshot. Apart from the genome-wide ePV (highlighted within the Structural variation track), no significant SNPs were detected across the three samples, confirming that the same strain was analyzed. MTase-related genes are annotated in dedicated tracks.

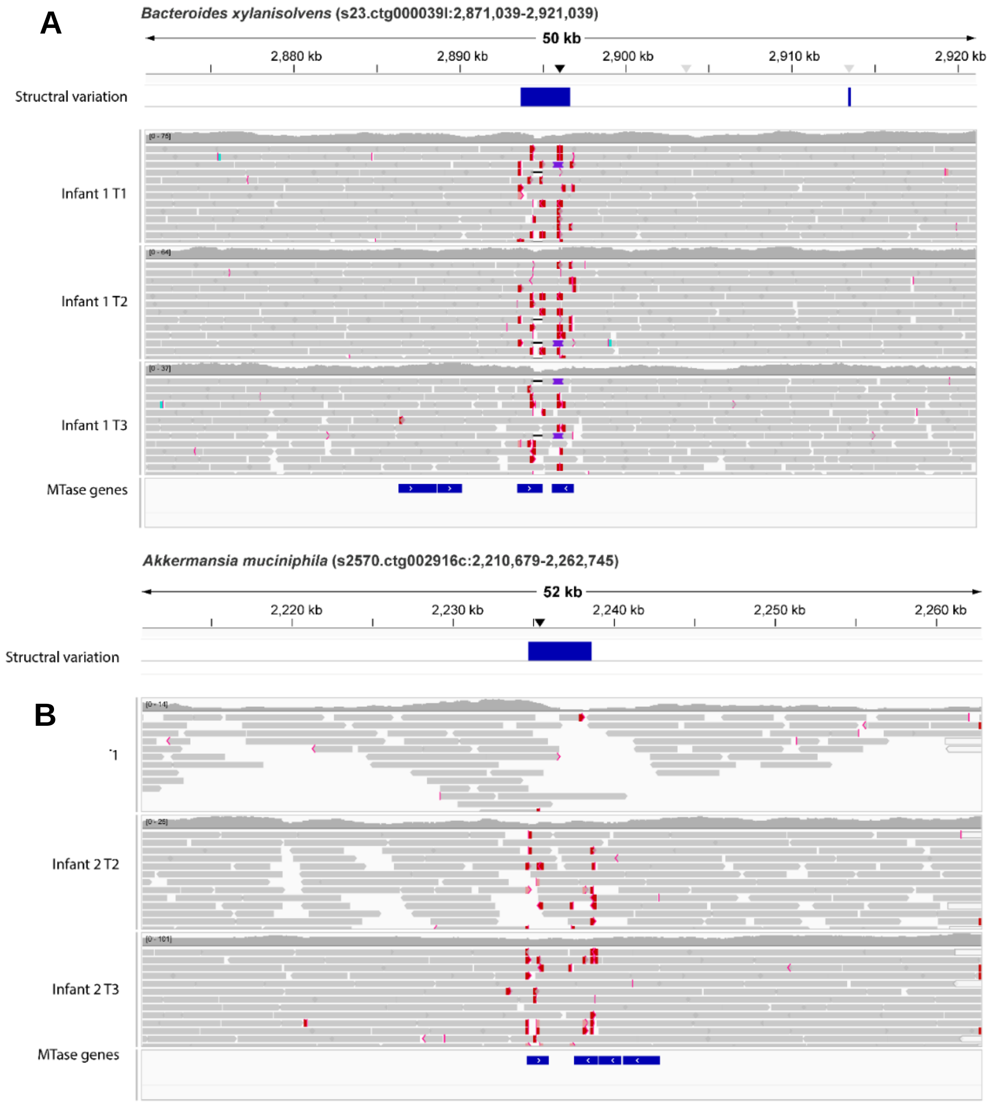

C

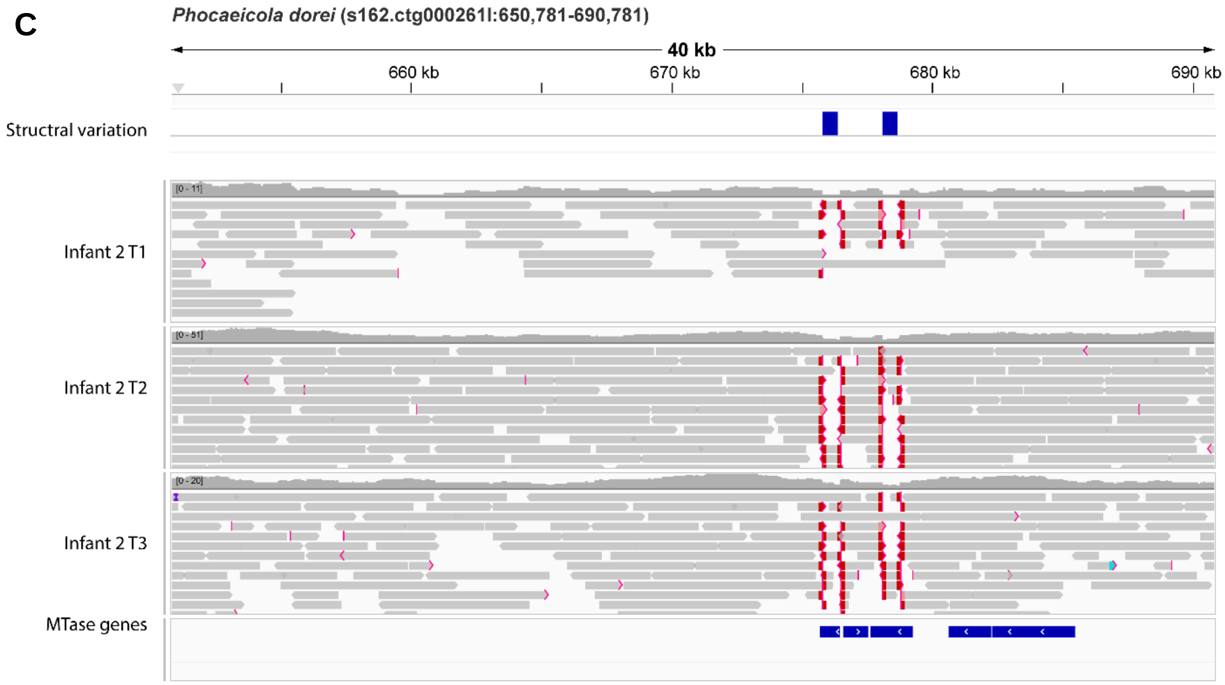
Phocaeicola dorei (s162.ctg000261l:650,781-690,781)
40 kb
660 kb
670 kb
680 kb
690 kb
Structral variation
Infant 2 T1
Infant 2 T2
Infant 2 T3
MTase genes

D

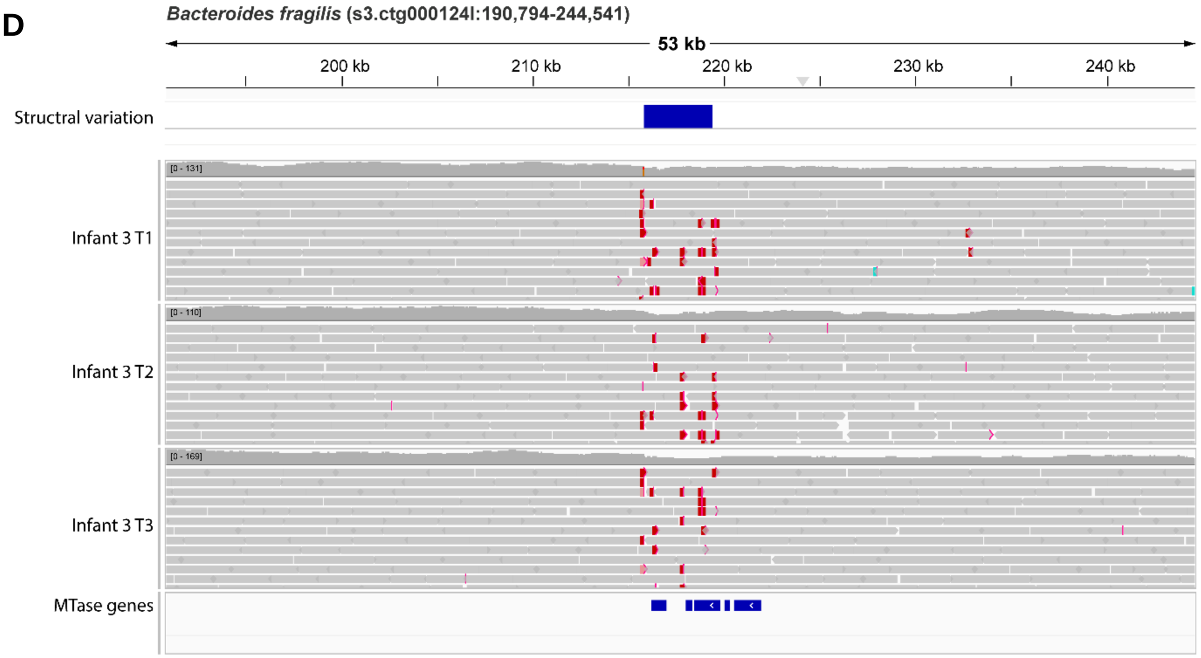
Bacteroides fragilis (s3.ctg000124l:190,794-244,541)
53 kb
200 kb
210 kb
220 kb
230 kb
240 kb
Structral variation
Infant 3 T1
Infant 3 T2
Infant 3 T3
MTase genes

E

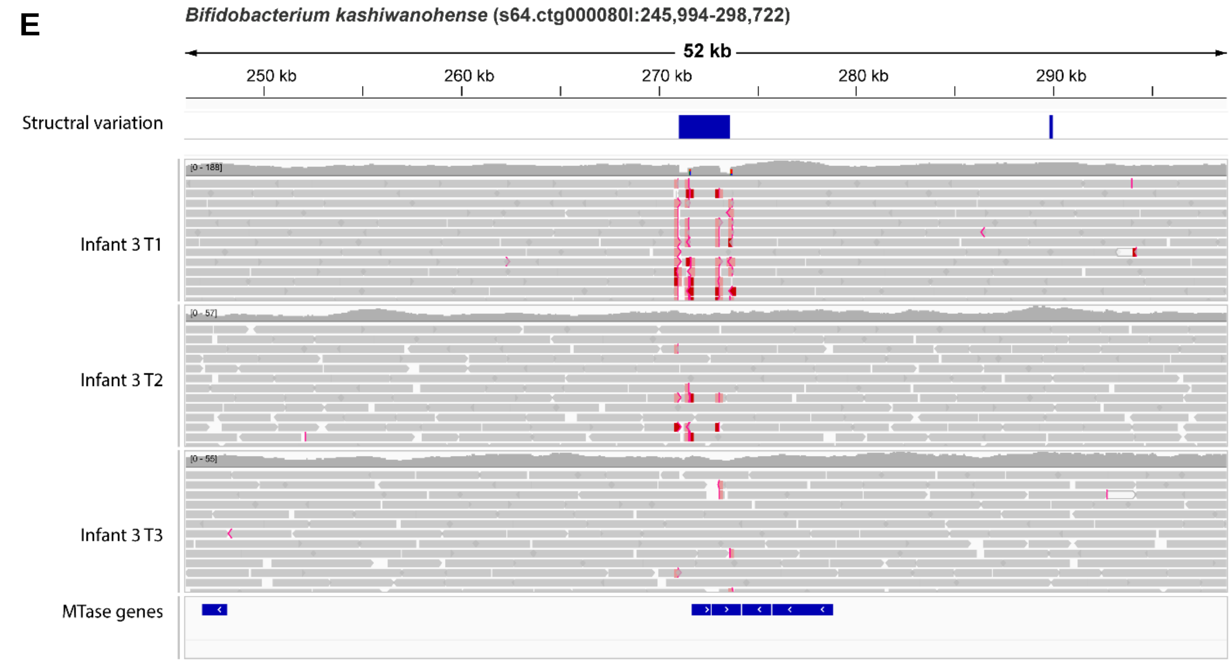
Bifidobacterium kashiwanohense (s64.ctg000080l:245,994-298,722)
52 kb
250 kb
260 kb
270 kb
280 kb
290 kb
Structral variation
Infant 3 T1
Infant 3 T2
Infant 3 T3
MTase genes

F

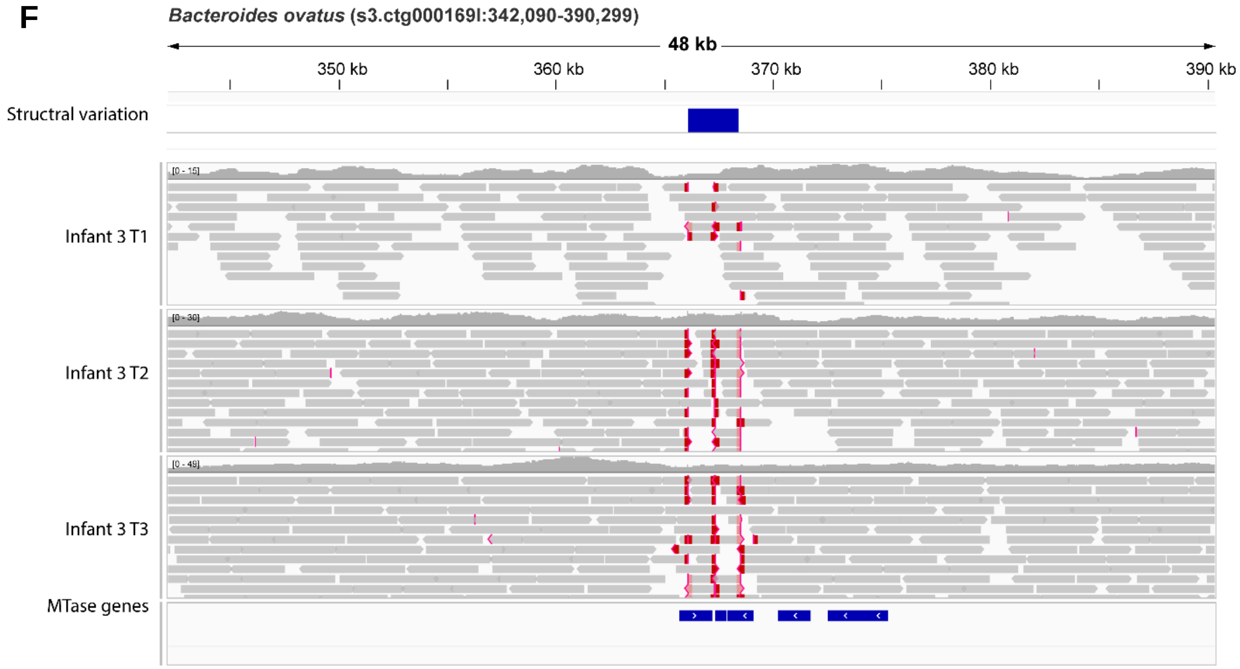
Bacteroides ovatus (s3.ctg000169l:342,090-390,299)
48 kb
350 kb
360 kb
370 kb
380 kb
390 kb
Structral variation
Infant 3 T1
Infant 3 T2
Infant 3 T3
MTase genes
